# Prenatal inflammation perturbs fetal hematopoietic development and causes persistent changes to postnatal immunity

**DOI:** 10.1101/2022.05.08.491095

**Authors:** April C. Apostol, Diego A. López, Eric J. Lebish, Clint H. Valencia, Mari Carmen Romero-Mulero, Polina Pavlovich, Gloria E. Hernandez, E. Camilla Forsberg, Nina Cabezas-Wallscheid, Anna E. Beaudin

## Abstract

Adult hematopoietic stem and progenitor cells (HSPCs) respond directly to inflammation and infection, resulting in both acute and persistent changes to quiescence, mobilization, and differentiation. Here we show that fetal HSPCs respond to prenatal inflammation *in utero*, and that the fetal response shapes postnatal hematopoiesis and immunity. Heterogenous fetal HSPCs showed divergent responses to maternal immune activation (MIA), including changes in quiescence, expansion, and lineage-biased output. Single cell transcriptomic analysis of fetal HSPCs in response to MIA revealed specific upregulation of inflammatory gene profiles in discrete, transient HSC populations, that propagated expansion of lymphoid-biased progenitors. Beyond fetal development, MIA caused the inappropriate expansion and persistence of fetal lymphoid-biased progenitors postnatally, concomitant with increased cellularity and hyperresponsiveness of fetal-derived innate-like lymphocytes. Our investigation demonstrates how inflammation *in utero* can direct the trajectory of output and function of fetal-derived immune cells by reshaping fetal HSC establishment.

## Introduction

The adult immune system responds rapidly to infection by increasing the production of immune cells that control infectious microbes (Schultze et al., 2019; Takizawa et al., 2012). This response includes the release of inflammatory cytokines, mobilization and activation of specific immune cells, and increased production of immune cells in the bone marrow (BM) needed to eradicate microbes. The ability of the immune system to initiate a sustained and systemic response to infection has recently been linked to the direct responsiveness of adult BM hematopoietic stem cells (HSCs) to inflammatory signals. Adult HSCs respond to an array of cytokines and TLR ligands *in situ* (Baldridge et al., 2010; Esplin et al., 2011; Essers et al., 2009; Furusawa et al., 2016; Nagai et al., 2006; Pietras et al., 2016; Schuettpelz and Link, 2013; Takizawa et al., 2017; Yamashita and Passegué, 2019), resulting in both acute and long-term changes to HSC function that ultimately shape the immune response.

In response to inflammation or infection, adult HSCs exit quiescence (Essers *et al*., 2009), show accentuated myeloid bias (Matatall et al., 2014; Pietras *et al*., 2016), and increased myeloid output, leading to the production of sufficient quantities of myeloid cells required to combat infection. Accumulating evidence suggests that in response to both normal and aging-related inflammation, myeloid-biased output is propagated by the disparate activation of myeloid-biased HSCs (Beerman et al., 2010; Haas et al., 2015; Matatall *et al*., 2014), as well as the downstream expansion of myeloid-biased multipotent progenitors at the expense of lymphoid progenitors (Pietras *et al*., 2016). Furthermore, persistent responsiveness of hematopoietic stem and progenitor cells (HSPCs) to inflammatory signals, programmed by epigenetic and metabolic alterations, has been suggested to underlie long-term functional changes to infection described in short-lived myeloid cells (Chavakis et al., 2019; Kaufmann et al., 2018; Mitroulis et al., 2018). These data suggest that inflammation can program long-term immune output from hematopoietic progenitors and has consequences for immune function.

Prenatal inflammation and infection shape the postnatal immune response, including susceptibility to infection, autoimmunity and hypersensitivity disorders, and response to vaccination (Apostol et al., 2020), but the drivers of persistent changes to offspring immunity are poorly understood. In the developing embryo, “sterile” inflammatory signaling is required for HSC emergence during early development (Espin-Palazon et al., 2018). However, beyond emergence, there is little information regarding whether developing HSCs respond *in utero* to maternal infection and inflammation, how they respond, and the consequences for the development of postnatal hematopoiesis and immunity. We hypothesize that the fetal HSC response to prenatal inflammation drives lasting changes to hematopoietic output in offspring that may help shape the postnatal immune response.

Here, we use a maternal immune activation (MIA) model to decipher the effects of *in utero* inflammation on fetal HSPCs, and examine the long-term impact to postnatal hematopoietic and immune system establishment. In direct contrast to myeloid bias invoked by inflammation in adult hematopoiesis, our *in vivo* analysis and single-cell transcriptional profiling reveal that prenatal inflammation evokes lymphoid bias by activating transient, lymphoid-biased fetal HSCs and propagating the expansion of lymphoid-biased HSPCs. In response to prenatal inflammation, the activation and inappropriate persistence of otherwise transient lymphoid-biased fetal HSCs leads to functional changes to postnatal hematopoiesis and immune output, including increased production and activity of fetal-derived innate-like lymphocytes. By demonstrating that heterogeneity of the fetal HSPC compartment underlies a differential response of prenatal hematopoiesis to inflammation, our findings have important implications for defining developmental origins of immune dysfunction.

## Results

### Prenatal inflammation induced by maternal immune activation (MIA) causes expansion of a developmentally-restricted HSC

To investigate how fetal hematopoietic progenitors respond to maternal inflammation, we employed a mouse model of maternal immune activation (MIA; (Meyer et al., 2009), where we mimicked a mild viral infection in pregnant mice with a single injection of the toll-like receptor 3 (TLR3) agonist Polyinosinic:polycytidylic acid, or pIC. We performed these experiments using the FlkSwitch mouse model (Boyer et al., 2011), which we previously used to identify a developmentally-restricted, lymphoid-biased hematopoietic stem cell (drHSC) with specific potential to produce fetal-restricted immune cells, including B1-B cells (Beaudin et al., 2016). In the FlkSwitch model, Flk2-driven expression of Cre results in a permanent genetic switch from Tomato (Tom) to GFP expression (Boyer et al., 2011). Both Tom+ and GFP+ HSCs are present during perinatal development (**Fig. 1A**), but only Tom+ HSCs persist and contribute functionally to the adult HSC compartment (Boyer et al., 2012; Boyer *et al*., 2011). We therefore denote **Tom+ HSCs** in the fetal liver (FL) as “conventional” HSCs, and GFP+ HSCs in the FL as “developmentally-restricted” HSCs, or **GFP+ drHSCs** (Beaudin et al., 2016).

**Figure 1.**
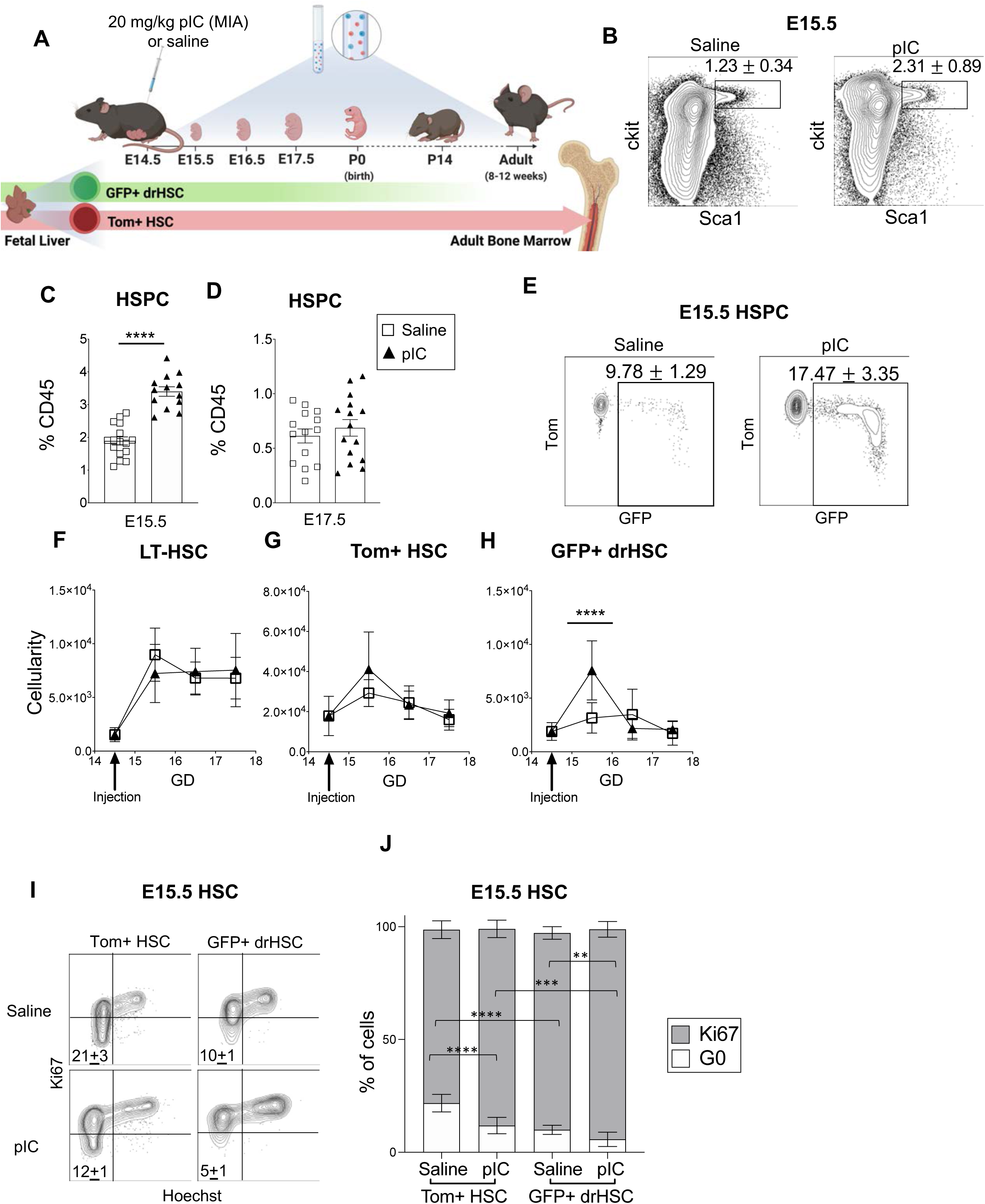
Fetal hematopoietic stem and progenitor cells (HSPCs) respond to prenatal inflammation induced by maternal immune activation (MIA) **A)** Timeline of MIA induction and analysis of fetal liver (FL) hematopoiesis. Pregnant WT dams mated to FlkSwitch males were intraperitoneal (i.p.) injected with 20 mg/kg Polyinosinic-polycytidylic acid (pIC) at embryonic day I 14.5. Both Tom+ HSCs and GFP+ drHSCs are present in the fetal liver (FL) during development, but only Tom+ HSCs persist into the adult bone marrow (BM). Image created with Biorender.com. **B)** Representative FACS plots of the hematopoietic stem and progenitor compartment “HSPC” (Live, Lin-, CD45+, cKit+, Sca1+) at E15.5, one day after pIC or saline induction. Numbers indicate average frequency ± SD. **C, D)** Frequency of HSPCs (as %CD45+ cells) in FL at E15.5 (C) and E17.5 (D). **E)** Representative FACS plots showing frequency of GFP+ cells within the HSPC compartment at E15.5 in mice described in Fig.1B. Data represent frequency ± SD. **F-H)** Quantification of long-term HSCs (**F)** (LT-HSCs; LSK Flk2-CD150+ CD48-), Tom+ HSCs (**G**; LSK CD150+ Tom+), and GFP+ drHSCs (**H;** LSK CD150+ GFP+) across gestation days (GD) following maternal injection of pIC or saline. For all above experiments, N=3-6 pups/litter representing at least 3 litters/condition. Data represent average ± SEM. *p ≤ 0.05; **p ≤ 0.01; ***p ≤ 0.001; ****p ≤ 0.0001. **I)** Representative FACS plots measuring proliferation by Ki67 expression and Hoechst staining of Tom+ HSC and GFP+ drHSCs after saline or pIC. N=3 litters/condition, 2-3 pups/litter. Data represent frequency ± SD. **J)** Quantification of Ki67 expression at E15.5 in both Tom+ HSCs and GFP+ drHSCs following pIC or saline in mice described in Fig. 1L. *p ≤ 0.05; ***p ≤ 0.001. Bars represent average ± SD.

To induce MIA, we injected pregnant C57BL/6J (WT) females crossed to FlkSwitch males with 20mg/kg pIC or saline control at embryonic day (E)14.5, when hematopoietic progenitors in the FL undergo massive expansion. We examined hematopoietic stem and progenitor cells (HSPCs) in offspring 1-3 days later, postnatally, and into adulthood (**Fig. 1A**). The dose of pIC used produced no obvious signs of illness in the mom. Crown-rump length (CRL) of E15.5 fetuses was lower in MIA litters compared to saline-treated litters **(SFig. 1A)** and onset of birth was delayed by one day in 35% of litters born (**SFig. 1B**). After birth, by postnatal day (P)14, there was a slight reduction in surviving pups from MIA **(SFig. 1C)** although those pups were on average, larger than controls **(SFig. 1D),** most likely due to longer gestation time (**SFig. 1B)**.

Clear changes to the fetal hematopoietic compartment were observed one day after MIA induction. HSPCs (CD45+, Lineage-, cKit+ Sca1+) showed a marked increase in frequency one day after injection (**Fig. 1B, C**), that was normalized by E17.5 (**Fig. 1D**). Despite increased HSPC frequency, HSPC cellularity at E15.5 was not significantly increased **(SFig. 1E)**, likely reflecting an overall decrease in CD45+ cells in the FL **(SFig. 1F)**. Importantly, as compared to adult BM HSPCs **(SFig. 2A-B)**, MIA did not induce Sca1 upregulation in the FL **(SFig. 2C),** including CD45+ cells **(SFig. 2D)** or long-term (LT)-HSCs (**SFig. 2E-F**), suggesting that increased fetal HSPC frequency was not caused by interferon-induced Sca1 upregulation. We also observed a significant increase in frequency of GFP-labeled cells across all HSPCs (**Fig. 1E**), consistent with expanded progenitors. Cellularity of LT-HSCs (CD45+ Lin-, cKit+, Sca1+ CD150+ CD48-; **Fig. 1F**), including Tom+ HSCs (CD45+, Lin-, cKit+, Sca1+, CD150+, Tom+; **Fig. 1G**) was unchanged from E15.5-E17.5 in response to MIA. In contrast, we observed a significant spike in the number of GFP+ drHSCs (CD45+, Lin-, cKit+, Sca1+, CD150+, GFP+) one day following MIA, after which cellularity returned to control levels by E16.5 (**Fig. 1H**).

To examine the mechanism underlying the specific expansion of the GFP+ drHSCs in response to MIA, we assessed proliferation at E15.5. MIA drove increased proliferation of both Tom+ HSCs and GFP+ drHSCs (**Fig. 1I-J**), but MIA drove virtually all GFP+ drHSCs out of G0 (**Fig. 1I-J**). Importantly, only increased proliferation of GFP+ drHSCs corresponded to a significant increase in cellularity one day later (**Fig. 1H**).

### Activation of a developmentally-restricted HSC drives downstream expansion of lymphoid-biased multipotent progenitors

As MIA induced a rapid response by HSCs that coincided with an increase in overall GFP expression across all HSPCs (**Fig. 1E**), we next examined how downstream hematopoietic progenitors responded to MIA, including short-term (ST)-HSCs and multipotent progenitor (MPP) subsets as defined phenotypically by Pietras et al (2015) (**Fig. 2A**). Cellularity of myeloid-biased ST-HSCs, MPP2, and MPP3 cells were unaffected by MIA (**Fig. 2B-D**), an observation that deviates from previously reported dynamics in the adult hematopoietic response to inflammation (Pietras, 2017). Similarly, committed myeloid progenitors were not expanded in response to MIA (**SFig. 1G, I**). In contrast, lymphoid-biased Flk2+ MPP4s were significantly expanded two-days post-MIA compared to saline treated controls (**Fig. 2E**) Expansion of Flk2+ MPP4s did not immediately translate into expansion of downstream committed lymphoid progenitors or mature lymphoid cells (**SFig. 1 H,J,K**), as expected based on differentiation dynamics of lymphoid lineages (Busch et al., 2015). Investigation of proliferation across all CD150-MPPs revealed increased proliferation at E15.5 and E16.5 (**Fig. 2F,G**), likely driving the significant increase in cellularity of MPP4s at E16.5. Expansion of both lymphoid-biased GFP+ drHSCs and lymphoid-biased Flk2+ MPP4s in response to MIA raised the possibility that prenatal inflammation specifically evoked a lymphoid-biased response in fetal progenitors.

**Figure 2.**
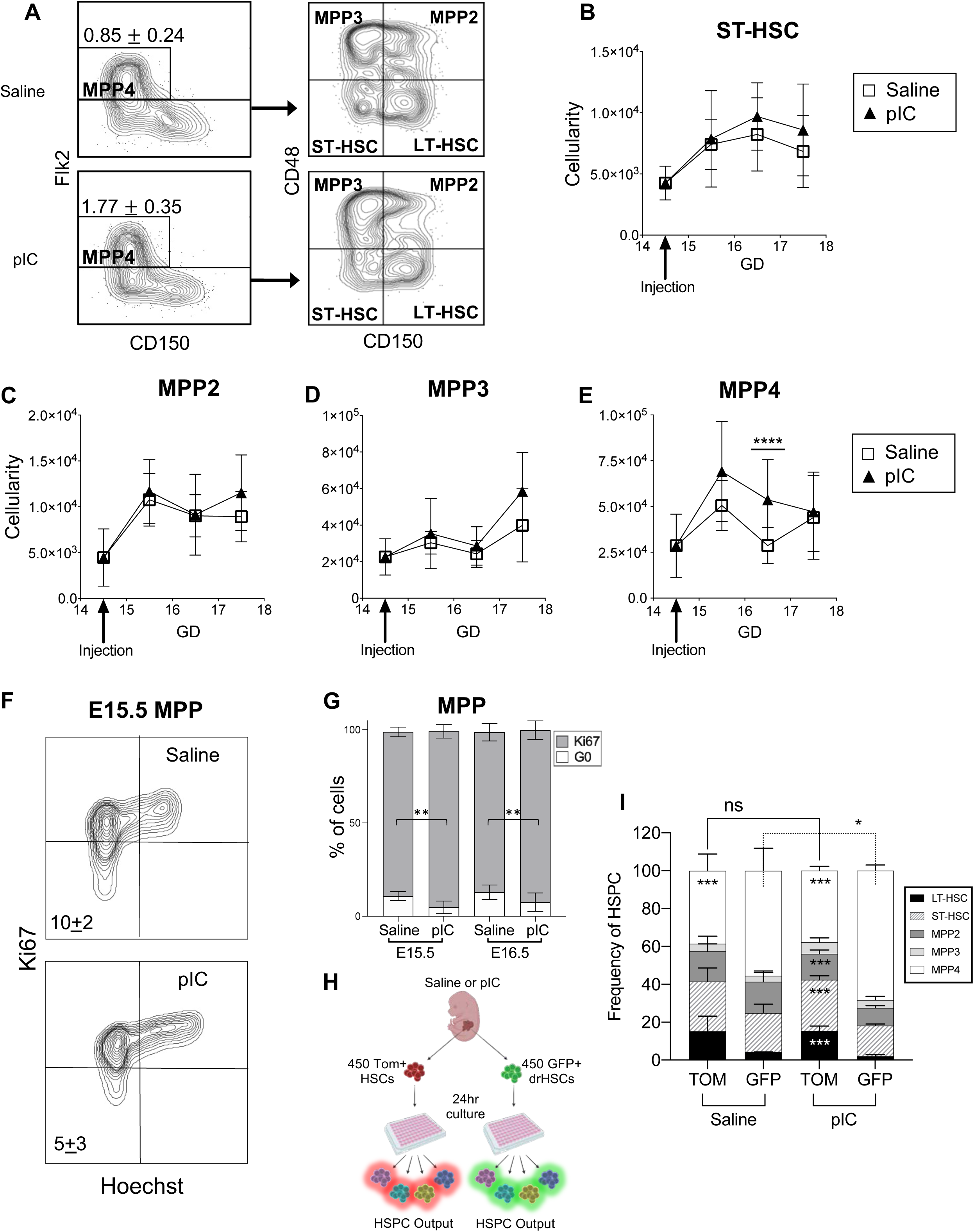
Lymphoid-biased fetal progenitors expand in response to prenatal inflammation induced by MIA. **A)** Representative FACS plots of the HSPC compartment (Live, Lin-, CD45+, cKit+, Sca1+) at E15.5. Numbers indicate average frequency ± SD of MPP4s in MIA and saline conditions. Data represent at least 3-6 pups/litter and 3 litters/condition. **B-E)** Cellularity of short-term HSCs (ST-HSC) **(B)**, MPP2s **(C)**, MPP3s (**D)**, and MPP4s **(E)** across gestation days (GD) 15-17 following MIA or saline. Data represent average ± SEM for at least 3-6 pups/litter and 3 litters/condition. ****p ≤ 0.0001. **F)** Representative flow cytometric plots of Ki67 expression and Hoechst staining in MPP (CD150-HSPC) populations after MIA or saline. Data represent average ± SD. N ≥ 3 litters/condition, 3-6 pups/litter. **G)** Quantification of Ki67 expression in MPPs (CD150-HSPC) from GD 15-17 after MIA or saline treatment. Data represent average ± SD for at least 3-6 pups/litter for 3 litters/condition at each timepoint. **p ≤ 0.01; ****p ≤ 0.0001. **H)** Schematic for *ex vivo* culture experiments. Two hours post *in vivo* treatment with pIC or saline, 450 Tom+ HSCs or GFP+ drHSCs were sorted and cultured. HSPC output was analyzed 24 hours later. **I)** Frequency of HSPC subsets generated from cultured saline or pIC treated Tom+ HSCs or GFP+ drHSCs. Data represent average ± SD from at least two independent experiments performed in triplicate. Significance indicated within bars represents statistically significant comparisons between TOM vs GFP within the same condition, and overhead significance brackets indicate statistically significant comparisons between same cell types across conditions. *p ≤ 0.05; *** p ≤ 0.001.

We previously demonstrated that the GFP+ drHSC is lymphoid-biased and lymphoid-lineage primed (Beaudin et al., 2016). To directly test whether expanded GFP+ drHSCs contributed to the increase in MPP4s, we isolated Tom+ HSCs and GFP+ drHSCs from FLs two hours post-saline or MIA induction and cultured them *ex vivo* over 24 hours and measured HSPC output (**Fig. 2H, SFig. 2L**). After 24 hours, both Tom+ and GFP+ drHSCs produced all HSPC subsets (**Fig. 2I**). GFP+ drHSCs generated significantly more MPP4s compared to Tom+ HSCs under saline conditions. Importantly, whereas MPP output from Tom+ HSCs was unchanged after MIA treatment, MPP4 output from GFP+ drHSCs was further exaggerated by MIA, at the expense of other MPP subsets (**Fig. 2I**).

### Prenatal inflammation induced by MIA affects self-renewal and hematopoietic output of developing HSCs

To investigate how MIA impacted function and output of Tom+ HSCs and GFP+ drHSCs, we performed transplantation assays and investigated long-term multi-lineage reconstitution (LTMR). We isolated Tom+ HSCs or GFP+ drHSCs from E15.5 FL one day post MIA or saline treatment, and competitively transplanted sorted cells with 5×10^5^ adult whole bone marrow (WBM) cells into lethally irradiated adult recipients (**Fig. 3A**). Recipient mice were monitored every 4 weeks for 16 weeks post-transplantation to assess mature blood lineage reconstitution in both primary and secondary transplants (**Fig. 3A**). Under control conditions, fewer recipients of GFP+ drHSCs were reconstituted as compared to recipients of Tom+ HSCs (**Fig. 3B**), and GFP+ drHSC recipients demonstrated lymphoid-biased contribution to peripheral blood (PB) output as compared to Tom+ HSC recipients **(SFig. 3A),** as previously reported (Beaudin, 2016). In primary transplantation, MIA did not impact LTMR frequency in recipients of either Tom+ HSCs or GFP+ drHSCs (**Fig. 3B**). MIA induced significantly increased granulocyte-monocyte (GM) output in PB (**Fig. 3C**) of primary Tom+ HSC recipients, although there were no differences in BM chimerism of HSPCs **(SFig. 3B-C),** myeloid progenitors, lymphoid progenitors, or mature cells after 18 weeks **(SFig. 3D, E)**. MIA did not affect PB (**Fig. 3C**) or BM progenitor output **(SFig. 3B-E)** in primary recipients of GFP+ drHSCs. In sharp contrast, secondary transplantation revealed a significant reduction in the number of Tom+ HSC recipients that exhibited LTMR (**Fig. 3D**), whereas the effect of MIA in secondary recipients of GFP+ drHSCs was less severe. Tom+ HSC exhaustion was evident in both PB output (**Fig. 3E**) and BM progenitor chimerism in MIA-treated Tom+ HSC recipients **(SFig. 3G-J**). In contrast, secondary recipients of GFP+ drHSCs did not exhibit any functional differences in response to MIA: both PB and BM chimerism was unaffected by MIA, and lymphoid-biased output was sustained (**Fig. 3E; SFig. 3F**). Together, serial transplantation assays revealed a disparate response of two fetal HSC populations, with an initial myeloid bias followed by subsequent loss of self-renewal potential in Tom+ HSCs, but not GFP+ drHSCs, following MIA.

**Figure 3.**
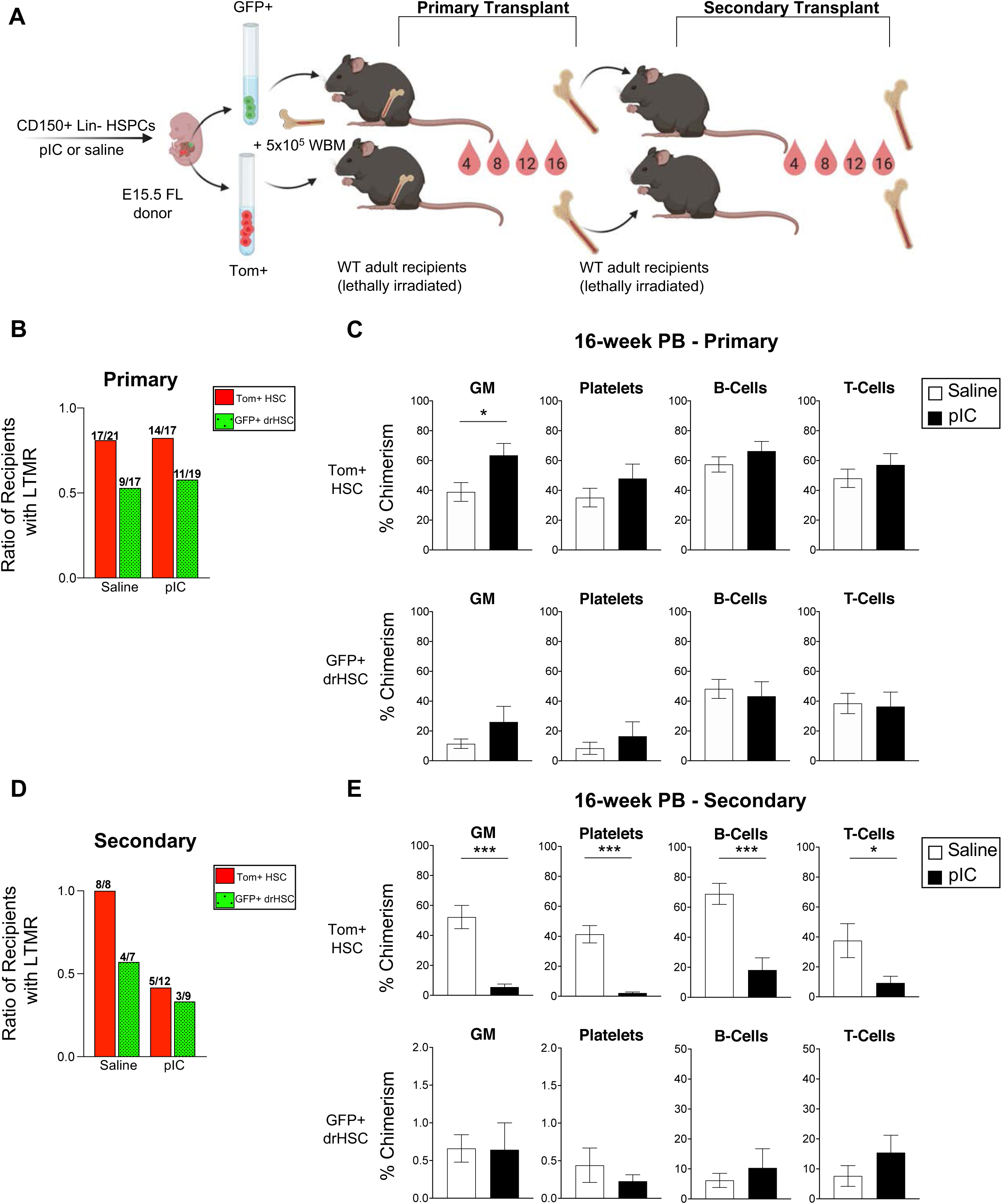
Prenatal inflammation induced by MIA causes persistent changes to fetal hematopoietic stem cells upon transplantation. **A)** Schematic of transplantation approach. After MIA at E14.5, E15.5 Tom+ HSCs or GFP+ drHSCs were sorted and transplanted into lethally irradiated C57BL/6 hosts (200 HSCs/host) along with 5×10^5^ whole bone marrow (WBM) cells. Peripheral blood (PB) was analyzed every 4 weeks until week 16. Image created with Biorender.com. **B)** The ratio of recipients with long-term multilineage reconstitution (LTMR) from transplants of Tom+ HSCs (red bar) and GFP+ drHSCs (patterned green bar) from pIC and saline-treated litters. N as indicated by numbers above bars, 13-19 recipients/condition. **C)** PB chimerism at 16-weeks of GM (granulocyte/macrophage), platelets (Plt), B-cells, and T-cells in recipients of Tom+ HSCs or GFP+ drHSCs, from same mice indicated in **B**. Data represent average ± SEM. *p ≤ 0.05. **D)** The ratio of recipients with long-term multilineage reconstitution (LTMR) from secondary transplants of 5×10^6^ WBM cells from primary recipients of Tom+ HSCs (red bar) and GFP+ drHSCs (patterned green bar) from mice indicated in **B**. N as indicated by numbers above bars, 7-12 recipients/condition. **E)** PB chimerism at 16-weeks of GM (granulocyte/macrophage), platelets (Plt), B-cells, and T-cells in secondary recipients described in **D**. Data represent average ± SEM. *p ≤ 0.05; ***p ≤ 0.001.

### Single-cell sequencing reveals the response of distinct fetal HSPCs to MIA

To further understand how MIA remodels hematopoietic development at the molecular level, we sorted E15.5 FL HSPCs from MIA or saline-treated dams and analyzed the transcriptional profiles of 23,505 cells using single-cell RNA sequencing (scRNA-seq). Unsupervised clustering analysis identified 14 distinct clusters (C0-C13) **(Fig. 4A, SFig. 4A).** Clusters segregated based on differential gene expression profiles and cell-cycle states (**SFig. 4B)**. MIA altered the distribution of cells across clusters (**Fig. 4B, C**). Differential response to MIA was observed among clusters: C1, C3, C5, C6, C9, and C10 were robustly expanded in response to MIA, whereas C2, C4, C7, and C8 were significantly reduced in response to MIA (**Fig. 4D**).

**Figure 4.**
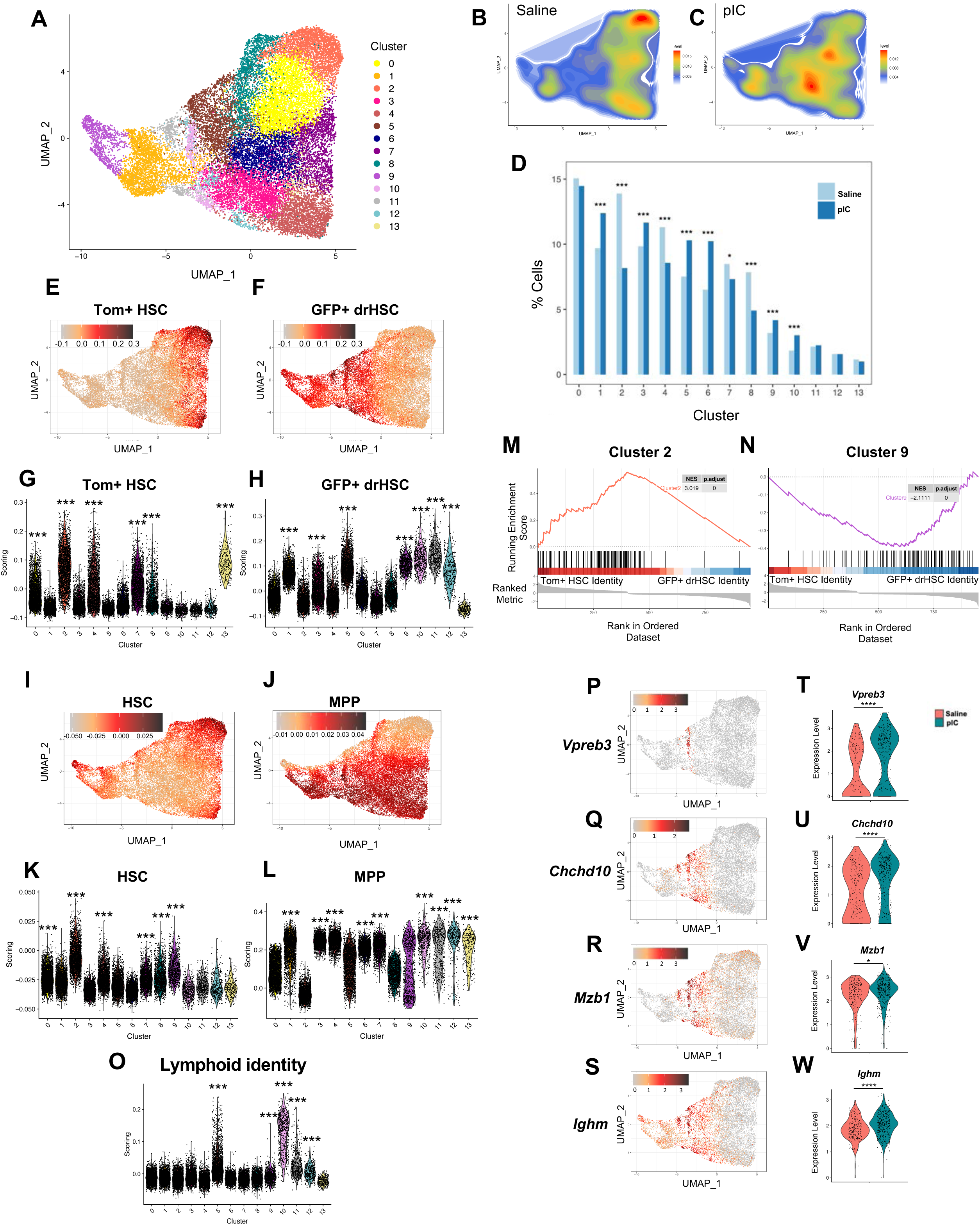
Prenatal inflammation induces distinct molecular changes in fetal HSCs. **A)** UMAP plot of single cell RNA-sequencing showing 14 identified clusters from sorted E15.5 FL Tom+ or GFP+ HSPCs (Lin-CD45+cKit+Sca1+) following pIC or saline treatment at E14.5. **B-C)** Representative density plots of clusters shown in **(A)** under saline **(B)** or pIC **(C)** conditions. **D)** Changes in percent of cells in each cluster, visualized in a barplot, in response to pIC or saline. Significance determined using Fisher tests. *p ≤ 0.05; ***p ≤ 0.001. **E, F)** UMAP plots showing enrichment scoring for TomISC **(E)** or GFP+ drHSC **(F)** signatures across clusters using reference bulk data set (Beaudin et al., 2016). Color bar indicates the enrichment scoring assessed by the AddModuleScore function. **G, H)** Violin plots showing enrichment scoring for Tom+ HSC **(G)** or GFP+ drHSC **(H)** signatures across clusters using a reference population data set (Beaudin et al., 2016). Wilcoxon signed rank test with respect to background. ***p ≤ 0.001. **I, J)** UMAP plots showing enrichment scoring for HSC **(I)** or MPP **(J)** signatures across clusters using reference bulk data set (Cabezas-Wallscheid et al., 2014). Color bar indicates the enrichment scoring assessed by the AddModuleScore function. **K, L)** Violin plots showing enrichment scoring for adult HSC **(K)** or MPP **(L)** signatures across clusters using a reference population data set (Cabezas-Wallscheid et al., 2014). Wilcoxon signed rank test with respect to background. ***p ≤ 0.001. **M, N)** Gene set enrichment analysis (GSEA) of Cluster C2 (M) and C9 (N) markers identified by FindMarkers function in a ranked set of differentially expressed genes between Tom+ HSC and GFP+ drHSCs (Beaudin et al., 2016). **(O)** Violin plots showing enrichment scoring for lymphoid gene signature across all clusters using a reference population data set (Cabezas-Wallscheid et al., 2014). Wilcoxon signed rank test with respect to background. ***p ≤ 0.001. **P-S)** UMAP plots showing gene expression levels of *Vpreb3* **(P)** *Chchd10* **I** *Mzb1* **(R)** and *Ighm* **(S)** under saline control conditions. **(T-Q)** Violin plots comparing expression of C10 markers under saline and pIC conditions. Wilcoxon signed rank test. *p ≤ 0.05; ****p ≤ 0.0001.

We utilized our previously published data set (Beaudin et al., 2016) to determine overlap of the phenotypical FL GFP+ drHSCs and Tom+ HSCs with our identified clusters. This resulted in the emergence of two divergent regions encompassing multiple clusters enriched for Tom+ HSC (**Fig. 4E, G**) or GFP+ drHSC signature (**Fig. 4F, H**). Surprisingly, several clusters that expanded in response to MIA significantly overlapped with the GFP+ drHSC signature (C1, C3, C5, C9, and C10), whereas clusters decreasing in response to MIA overlapped with the Tom+ HSC signature (C2, C4, C7, and C8). TdTomato and eGFP transcript expression was also enriched in clusters overlapping with Tom+ HSC and GFP+ drHSC signatures, respectively **(SFig. 4C),** although overlap was not precise due to broad GFP expression across MPPs within the FlkSwitch model (Boyer *et al*., 2011).

To annotate clusters responding to MIA, we determined HSC and MPP identity based on adult transcriptional profiles from previously published datasets (Cabezas-Wallscheid et al., 2014). Clusters segregated into either HSC- (**Fig. 4I, K**) or MPP-like populations (**Fig. 4J, L**). Two clusters, C2 and C9, were the most enriched for HSC identity and showed no significant enrichment for MPP genes. Gene set enrichment analysis (GSEA) of HSC-like C2 revealed best concordance with Tom+ HSC identity (**Fig. 4M**) and was defined quiescent cell cycle state (**SFig. 4B**) and by expression of canonical HSC genes, including *Mecom* **(SFig. 4A)**, *Hlf* **(SFig 4A),** *Mllt3* **(SFig. 4D),** and *Meis1* **(SFig 4E)** (Gazit et al., 2013). The remaining Tom+ HSC clusters -- C0, C4, C7, C8 and C13 (**Fig. 4G**) -- overlapped with both HSC and MPP signatures (**Fig. 4I-L**) and showed significant enrichment of megakaryocyte-associated genes **(SFig. 4F)** and no enrichment for lymphoid- (**Fig. 4O**) or myeloid-associated genes **(SFig. 4G)**. The identification of HSC-like C2 within the clusters most enriched for Tom+ HSC identity is consistent with HSC activity within the Tom+ HSC fraction.

Clusters enriched for the GFP+ drHSC signature -- C1, C3, C5, C9 10, C11, and C12 (**Fig. 4H**) -- exhibited varying degrees of stemness (**Fig. 4I-L**), lymphoid- (**Fig. 4O**) and myeloid-priming **(SFig. 4G)**. Uniquely, C9 was specifically enriched for both GFP+ drHSC signature (**Fig. 4F, H**) and adult HSC signature (**Fig. 4K**). Additionally, C9 was the only cluster to be significantly enriched for lymphoid-, myeloid-, and megakaryocyte-associated genes (**Fig. 4O**, **SFig 4F-G**). Although C9 did not express the same set of canonical HSC genes expressed by C2, C9 expressed other canonical and non-canonical HSC markers, including HLA genes **(SFig. 4A)**, *Spi1* (Staber et al., 2013) **(SFig. 4H)**, *Pbx1* (Ficara et al., 2008) **(SFig. 4I)**, *Cd74* (Luckey et al., 2006) **(SFig. 4A, J)**, and Tet2 (Moran-Crusio et al., 2011) **(SFig. 4K)**. C9 was heterogenous for cell cycle markers **(SFig. 4B),** with a subset of cells appearing highly quiescent. C5, C9, C10, C11, and C12 were all highly enriched for lymphoid gene signature (**Fig. 4O**) while C1, C5, C9, C11, and C12 were all enriched for myeloid gene signature **(SFig 4G)**. Notably, C10 expressed genes important for early B-cell and innate-like B-cell development and identity including *Vpreb1/3, Ccr9, Ebf1, Ighm, Chchd10,* and *Mzb1* (**Fig. 4P-S**) and was particularly enriched for the GFP+ drHSC signature (**Fig. 4H**). Furthermore, MIA accentuated lymphoid lineage priming in C10, as demonstrated by significant upregulation of the same lymphoid-associated genes (**Fig. 4T-W**). Importantly, overlap of multiple stem cell-, lymphoid-, and myeloid-associated clusters with the phenotypically-defined GFP+ drHSC reflects both multipotency and lymphoid bias of the GFP+ drHSC *in vivo*.

### Fetal HSPCs respond distinctly to inflammatory cytokines induced by MIA

*In utero,* the fetus may be passively exposed to maternal cytokines or other inflammatory mediators through the placental barrier or alternatively, the response of placental or fetal cells to maternal inflammation may translate into a discrete fetal immune response (Apostol *et al*., 2020). To interrogate potential signaling mechanisms that drive the specific response of lymphoid-biased FL HSPCs, we examined and compared inflammatory cytokine profiles in maternal serum, amniotic fluid (AF) and FL supernatant one day post MIA, at E15.5. IFN-ɑ and IL-23 were significantly increased in maternal serum (**Fig. 5A, SFig. 2G**). In AF, we observed an increase in levels of IFN-ɑ, IFN-β, IL-27, and GM-CSF in response to MIA (**Fig. 5B, C**). In contrast, the FL only exhibited upregulation of two cytokines: IL-1ɑ and IFN-ɑ (**Fig. 5D, E**). The increase in type I interferons in the maternal serum, AF, and FL suggest that type I interferon-mediated inflammation is evoked from either maternal sources of cytokine **(Fig. 5A, SFig. 2G),** or non-HSPC populations in FL, as IFN-α, IFN-β, and IL-1ɑ cytokine was not produced directly from HSPCs (data not shown). HSPCs also did not express TLR3 **(SFig. 2H)**, suggesting they did not respond directly to pIC.

**Figure 5.**
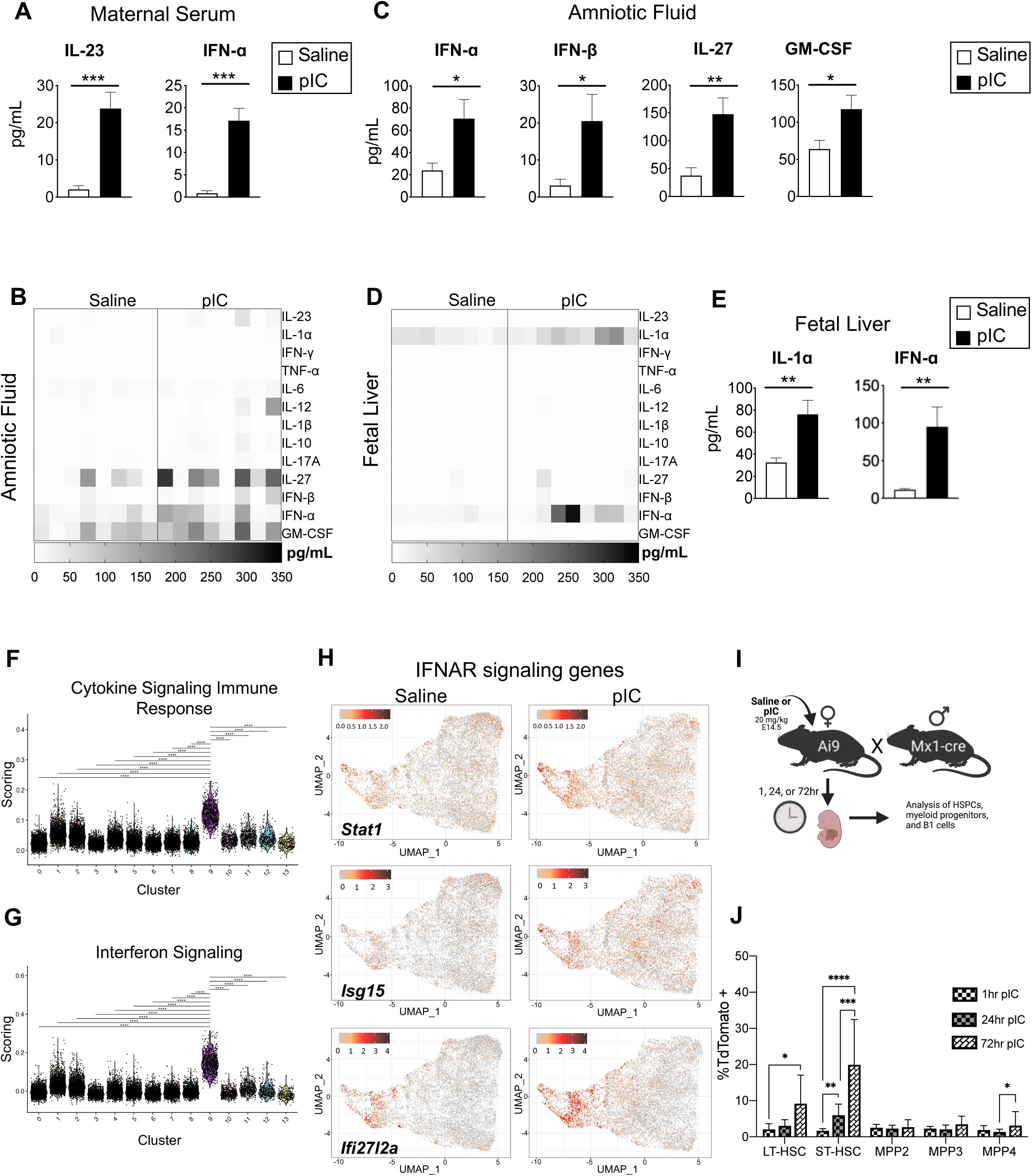
MIA-induced inflammatory cytokines specifically activate fetal HSPCs. **A)** Absolute quantification of cytokines in maternal serum at E15.5 as measured by LegendPlex assay. Data represent average ± SD. ***p ≤ 0.001. **B)** Heatmap displaying cytokines measured in amniotic fluid collected from E15.5 fetuses in litters following pIC or saline. **C)** Absolute quantification of significantly upregulated cytokines in amniotic fluid at E15.5. Data represent average ± SD. *p ≤ 0.05; **p ≤ 0.01. **D)** Heatmap displaying cytokines in fetal liver supernatant at E15.5 following pIC or saline. **E)** Absolute quantification of significantly upregulated cytokines in fetal liver supernatant at E15.5. For all experiments, N= 3 litters/condition with 3 pups/litter. Data represent average ± SD. **p ≤ 0.01. **F)** Single cell RNA-sequencing clusters scored against database of pathways (reactome.org) integral to cytokine signaling in the immune system. Scoring indicates enrichment assessed by the AddModuleScore function within individual clusters. Wilcoxon signed rank test. ****p <0.0001. **G)** Single cell RNA-sequencing clusters scored against (reactome.org) database of pathways integral to interferon signaling in the immune system. Scoring indicates enrichment assessed by the AddModuleScore function within individual clusters. Wilcoxon signed rank test. ****p <0.0001. **H)** UMAP plots showing expression levels of genes associated with interferon alpha receptor (IFNAR) signaling following treatment with pIC or saline. Color bar indicates the relative expression level. **I)** Schematic of Mx1 Cre lineage-tracing experiments designed to determine the immediate response of fetal HSPCs to pIC-induced prenatal inflammation. Mx1-Cre males were crossed to Ai9 reporter females, and pregnant females were treated with 20mg/kg pIC or saline at E14.5. **J)** Frequency of cells labeled by Tom+ fluorescence in fetal liver HSPC populations 1-, 24-, and 72-hours following induction of Mx1 Cre by prenatal pIC or saline. Frequency of label (%TdTomato+) was determined by taking the average background labeling in saline-treated controls and subtracting it from the frequency of label in pIC-treated mice. N=6-14 representative of at least two independent experiments. Data represent average ± SD. *p ≤ 0.05, **p< 0.01, ***p ≤ 0.001, ****p ≤ 0.0001.

To gain additional insight into possible interactions between fetal HSPCs and cytokines observed in the fetal microenvironment, we scored the enrichment of immune-related cytokine signaling gene signatures in our single-cell sequencing data. Only C9, a stem cell-like cluster that overlapped significantly with a GFP+ drHSC expression signature, was significantly enriched for expression of genes associated with immune cytokine signaling, as compared to all other clusters (**Fig. 5F**). Further examination of IL-1α and IFN-α signaling -- cytokines upregulated in the FL following MIA (**Fig. 5D,E**) -- revealed significant enrichment for interferon signaling pathways within C9 (**Fig. 5G**). In particular, interferon-α/β receptor (IFNAR) pathway genes *Stat1*, *Isg15*, and *Ifi27l2a* were highly expressed within C9 and increased expression following MIA (**Fig. 6H**). In contrast, overall interleukin-1 signaling expression was unchanged or decreased across clusters following MIA **(SFig. 2I),** suggesting that IFN-α signaling is likely a major driver of the inflammatory response within fetal HSPCs.

**Figure 6.**
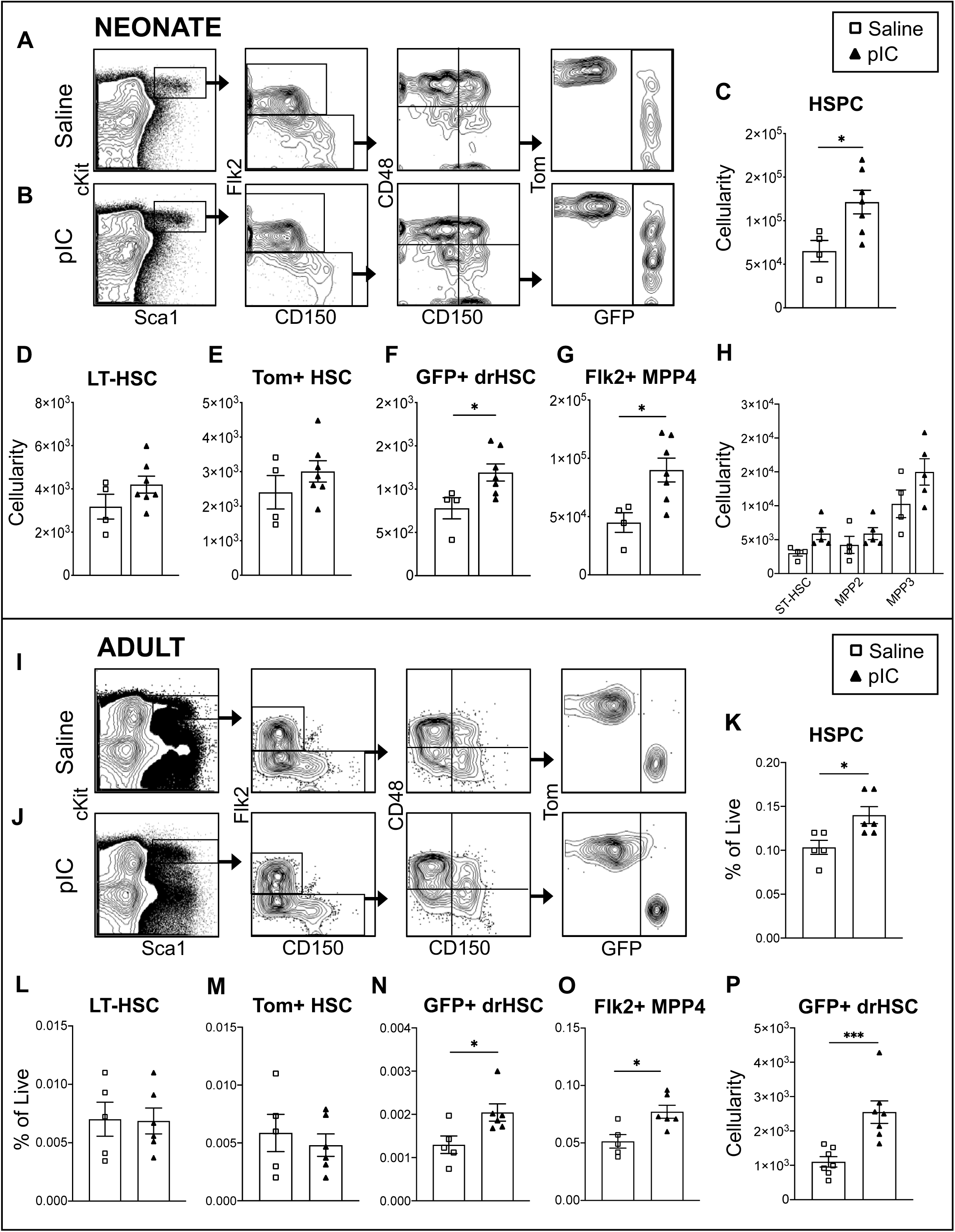
Prenatal inflammation induced by MIA affects postnatal hematopoiesis. **A, B)** Representative FACS plots indicating gating of the bone marrow HSPC compartment in saline **(A)** or pIC-treated **(B)** offspring at postnatal day (P)14. Tom+ HSC and GFP+ drHSC are denoted by fluorescence within the LT-HSC (CD150+CD48-Flk2-HSPC) compartment in FlkSwitch mice. **C-H)** Absolute quantification of HSPCs (gated as shown in Fig. 2A) including total HSPCs **(C)**, LT-HSCs (**D**), Tom+ HSCs **(E)**, GFP+ drHSCs **(F)**, Flk2+ MPP4s **(G)**, and ST-HSCs, MPP2s, and MPP3s **(H)**, gated as shown in A. N= 4-7 pups per litter, for at least three litters per condition. Data represent average ± SEM. *p ≤ 0.05. **I, J**) Representative flow cytometric analysis of BM HSPC compartment at adulthood (8-10 weeks) from saline **(I)** or pIC-treated **(J)** litters. **K-O)** Frequency of adult BM HSPCs (gated as shown in Fig. 2A) including total HSPCs **(K)**, Total LT-HSCs (**L**), Tom+ HSCs **(M)**, GFP+ drHSCs **(N)**, and Flk2+ MPP4 **(O)** (gated as shown in A). *p ≤ 0.05. Frequency of additional HSPCs can be found in **SFig. 5F**. **P**) Absolute quantification of GFP+ drHSCs. Absolute quantification of additional HSPCs can be found in **SFig. 5A-E**. N=6 representing at least three separate litters/experiments per condition. Data represent average ± SEM. ***p ≤ 0.001.

To further elucidate the response of fetal HSPCs to type I interferons, we crossed mice expressing Cre recombinase under control of the IFN-inducible Mx1 promoter (Mx1^Cre^) to an Ai9 reporter mouse (R26^LSL-tdTomato^) and ascertained labeling of fetal HSPCs 1-, 24-, and 72-hours following pIC or saline control (**Fig. 5I**). Mx1-Cre labeling at 1 hour in response to MIA was evident across all HSPCs via Tomato (Tom+) expression (**Fig. 5J**). By 24 hours post MIA, labeling was significantly enriched in ST-HSCs. After 72 hours, ST-HSCs exhibited the highest degree of labeling, and LT-HSCs also showed significant increased frequency of labeling. MPP4s similarly exhibited a significant increase in frequency of labeling 72 hours post MIA, whereas labeling of MPP2 and MPP3 subsets did not significantly increase over time (**Fig. 5J**). Analysis of Mx1-Cre labeling within myeloid progenitors (cKit+ Sca1-CD45+ Lin-) showed minimal labeling 24 hours post MIA **(SFig. 2J)**. Similarly, B1 B-cells displayed little to no Tom+ expression **(SFig. 2K)**. As activation of the Mx1 promotor is stringent and transient, even under continued exposure to type-1 interferons in culture (Staeheli et al., 1986), the increase in frequency of labeling in ST-HSCs over time reflects expansion of labeled cells after pIC induction, rather than *de novo* Mx1-Cre labeling. Because FL GFP+ drHSCs are enriched in the CD150^mid^ LSK fraction in FlkSwitch mice, the increased label of ST-HSCs in Mx1^Cre^Ai9 mice likely reflects combinatory labelling of both GFP+ drHSCs and ST-HSCs, the former demonstrating greater proliferation and cellularity 24 hours post MIA (**Fig. 5H-J**). Although ST-HSCs exhibited the greatest increase in label 24-72 hours post MIA, we observed no significant increase in ST-HSC cellularity *in vivo* (**Fig. 2B**), whereas our *in vivo* observation reveal that GFP+ drHSCs are expanded one day after MIA induction, followed by significantly increased MPP4 cellularity two days post MIA. Together, these data suggest that the response to MIA is driven by an immediate HSC response to type-1 interferons.

### Developmental effects of prenatal inflammation on hematopoiesis persist postnatally

As MIA perturbed fetal hematopoiesis and fetal HSPC function, we next sought to determine if MIA affected hematopoiesis in the postnatal period. Approximately three weeks after MIA induction, at postnatal day (P) 14, we observed sustained quantitative changes to the HSPC compartment (**Fig. 6**). The HSPC compartment (Lin-CD45+cKit+Sca1+; **Fig. 6A-B**) was significantly expanded in frequency and number in offspring in response to MIA (**Fig. 6B-C**). Although the overall LT-HSC compartment was unchanged in response to MIA (**Fig. 6D**), demarcation of the LT-HSC compartment into Tom+ HSCs and GFP+ drHSCs revealed specific expansion of the GFP+ drHSC (**Fig. 6F**) in the absence of changes to the Tom+ HSC compartment (**Fig. 6E**). In terms of absolute cellularity, expansion of the entire HSPC compartment could be attributed almost entirely to a 2-fold expansion of Flk2+ MPP4s (**Fig. 6G**), whereas other MPP subsets were unchanged in the postnatal period (**Fig. 6H**). This is the precise pattern we observed in response to MIA during the fetal period, suggesting a lasting effect of MIA on HSPCs.

As MIA perturbed hematopoiesis in offspring into the postnatal period, we examined if prenatal inflammation could drive lasting changes to adult hematopoiesis (**Fig. 6I-P; SFig. 5A-F**). Examination of the BM HSPC compartment in adult offspring following MIA revealed sustained expansion of HSPC frequency (**Fig. 6I-K**). As observed in the early postnatal period, (**Fig. 6A-H**), frequency of adult lymphoid-biased MPP4 subset (**Fig. 6O**) was increased in response to MIA, while cellularity and frequency of all other MPP subsets were not significantly affected (**SFig. 5E, F**). Frequency (**Fig. 6L, M**) and cellularity (**SFig. 5B, C**) of LT-HSCs and Tom+ HSCs were also unchanged in response to MIA. Although we have previously demonstrated both phenotypically (Boyer *et al*., 2011) and functionally (Boyer *et al*., 2012) that GFP+ drHSCs are virtually absent within the adult HSC compartment, we observed the presence of GFP+ cells within the phenotypic LT-HSC compartment in saline-treated adults (**Fig. 6N,P**). Nonetheless, we still observed sustained expansion of GFP+ drHSCs into the adult BM by both frequency (**Fig. 6N**) and cellularity (**Fig. 6P**) as compared to saline-treated controls. These data suggest that prenatal inflammation during fetal development imparts specific and lifelong changes to HSPC cellularity that can be traced well into adulthood.

### Persistent changes to hematopoiesis induced by prenatal inflammation affect innate-like immune cell function

We have previously shown that the GFP+ drHSC specifically generates fetal-derived innate-like lymphocytes during development, including B1 B-cells and MZ B-cells (Beaudin *et al*., 2016). As MIA caused persistent changes to the developing HSC compartment, including the specific expansion and inappropriate persistence of the GFP+ drHSC postnatally, we hypothesized that the inappropriate expansion of the GFP+ drHSC would subsequently affect the establishment of their innate-like immune cell progeny.

To determine the effect on innate-like lymphocyte establishment, we enumerated innate-like B-cells in the peritoneal cavity (PerC) and marginal zone (MZ) B-cells in the spleen in offspring of MIA- and control-treated litters at postnatal day (P) 14 (**Fig. 7A-C**). Whereas conventional B2 cells were unchanged in response to MIA, both CD5+ B1-a cells and CD5-B1-b cells were significantly expanded in P14 offspring following MIA (**Fig. 7A**). Similarly, cellularity of MZ splenic B-cells was increased in MIA offspring (**Fig. 7B**), whereas conventional follicular B2 cells were unchanged (**Fig. 7C**). Notably, fetal B cells did not expand during the fetal period in response to MIA (**SFig. 1H,K**) and were not interferon-responsive, as determined by negligible Mx1-Cre labeling (**SFig. 2K**), suggesting that postnatal expansion was due directly to an expanded GFP+ drHSC precursor. Specific expansion of fetal-derived innate-like lymphocyte progeny coincident with the postnatal expansion of the GFP+ drHSC precursor suggests that sustained changes to the HSC compartment led to measurable differences in fetal-derived innate-like immune cells postnatally.

**Figure 7.**
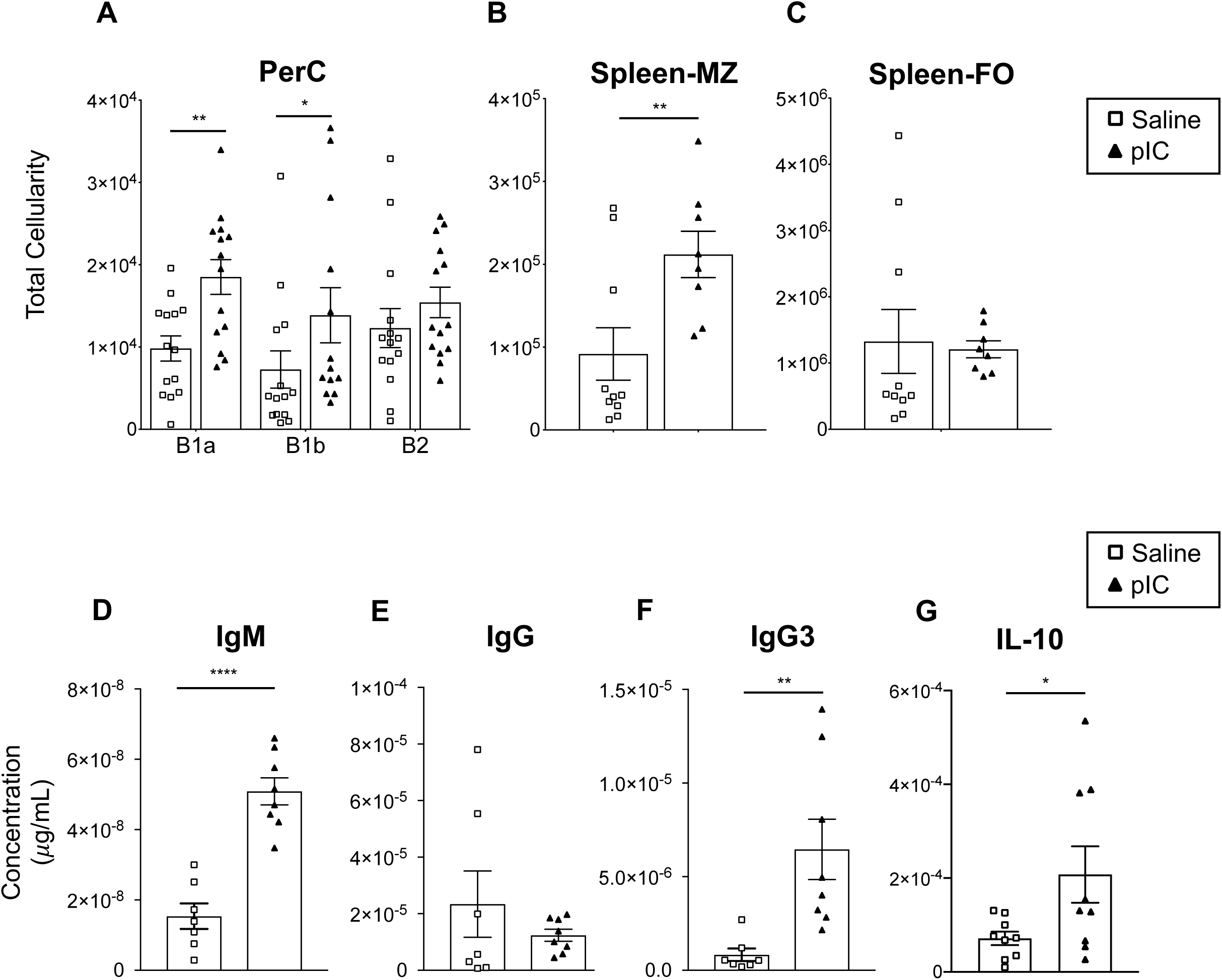
Prenatal inflammation induced by MIA shapes the function and output of fetal-derived immune cells. **A)** Absolute quantification of peritoneal IgM+ B-cell subsets, B1-a (CD5+CD11b+), B1-b (CD5-CD11b-) and B2 (CD5-CD11b-) cells in P14 offspring following pIC or saline at E14.5. **B,C)** Absolute quantification of marginal zone (MZ; CD21+CD23-) **(B)** and follicular zone (FZ; CD21-CD23+) **(C)** IgM+ B-cells in the spleens of P14 offspring following pIC or saline treatment at E14.5. **D-F)** Quantification of peritoneal IgM **(D)**, total IgG **(E)**, and IgG3 **(F)** by ELISA normalized for cellularity in pIC- and saline-treated offspring at P14. **G)** Quantification of IL-10 production following stimulation with LPS and PMA of equivalent numbers of peritoneal B-1 cells isolated from P21 offspring following pIC- or saline treatment at E14.5. N = 7-14 representing a least three litters per condition. Data represent average ± SEM. *p<0.05, **p<0.01, ****p<0.0001.

B1 B-cells function as a first line of immunity in neonates by producing “natural” antibodies (Baumgarth, 2011; Montecino-Rodriguez and Dorshkind, 2006). To define whether observed changes in innate-like lymphocyte cellularity also translated into changes in postnatal cellular function, we profiled natural antibody production in offspring following MIA (**Fig. 7D-F**). Coincident with expanded B1-B cell cellularity, we observed significantly more natural antibody, including IgM (**Fig. 7D**) and IgG3 (**Fig. 7F**) in the PerC of MIA-treated offspring, whereas total IgG was unchanged (**Fig. 7E**). Importantly, these results were normalized for cell number, suggesting that B1-B cells from MIA-treated offspring are not only expanded in number, but also produce more natural antibody on a per-cell basis as compared to saline controls. Furthermore, B1-B cells from MIA-treated offspring also produced more IL-10 upon stimulation (**Fig. 7G**), suggesting a greater propensity to respond to secondary immune challenge. These data suggest that the early hematopoietic response to *in utero* inflammation not only reshapes HSC function and establishment, but also alters output and function of developmentally regulated immune cells.

## Discussion

Adult HSCs respond directly to inflammation, resulting in both acute and sustained changes to HSC function that drive the systemic immune response to infection (Pietras, 2017). Whereas in the adult BM, HSCs are directly exposed to pathogens, cytokines, and other inflammatory mediators through circulation, exposure of the fetus to prenatal inflammation is necessarily limited by the maternal-fetal barrier (Ander et al., 2019). Maternal infection and microbial exposure during pregnancy has been linked to altered immune outcomes in infants (Apostol *et al*., 2020), but any causal relationship has not been clearly established, and there is little information regarding how *in utero* inflammation might be interpreted by the developing fetal immune system. “Sterile” inflammatory signaling is required for HSC emergence and specification during early development (Espin-Palazon *et al*., 2018; He et al., 2015), but to our knowledge there has been no direct investigation of how *in utero* inflammation directly influences fetal hematopoiesis. Here we demonstrate, for the first time, that developing fetal HSCs respond directly to *in utero* inflammation. To test the fetal HSC response to inflammation, we used a mouse model of maternal immune activation (MIA), in which injection of pIC at mid-gestation mimics an immune response to viral infection during pregnancy (Meyer *et al*., 2009). Importantly, the use of MIA allowed us to distill the effect of *in utero* inflammation in the absence of overt fetal infection. We also performed these experiments using our “FlkSwitch” model (Beaudin *et al*., 2016; Boyer *et al*., 2011) in which we have previously described two distinct fetal progenitors: a Tom+ HSC that persists into adult BM, and a lymphoid-biased, GFP+ developmentally-restricted hematopoietic stem cell (drHSC) that is transient and exists only in the perinatal period. This platform allowed us to test the response of a dynamic and heterogeneous fetal hematopoietic compartment to inflammation *in utero*. Our investigation reveals that prenatal inflammation directly influences fetal hematopoiesis and drives both an immediate and sustained response in otherwise transient fetal progenitors, thereby affecting the output and function of fetal-derived immune cells.

In order to respond appropriately to injury and inflammation, adult hematopoietic progenitors exhibit myeloid bias and increased myeloid output in response to a range of inflammatory cues (Haas *et al*., 2015; Matatall *et al*., 2014; Pietras *et al*., 2016). Myeloid bias is propagated both by the specific activation of myeloid-biased HSCs to inflammatory cues (Beerman *et al*., 2010; Gekas and Graf, 2013; Haas *et al*., 2015; Mann et al., 2018), as well as through downstream expansion of myeloid-biased progenitors, including MPP2 and MPP3, at the expense of lymphoid-biased MPP4s (Pietras *et al*., 2016). In sharp contrast, our analysis of the response of fetal hematopoiesis to prenatal inflammation *in situ* reveals that inflammation during the fetal period invokes a lymphoid-biased response. Using the FlkSwitch model, our analysis reveals that prenatal inflammation activates lymphoid-biased fetal GFP+ drHSCs and drives the downstream expansion of lymphoid-biased MPPs, including Flk2+ MPP4s. *Ex vivo* culture experiments confirmed that GFP+ drHSCs preferentially generate MPP4s, and that this bias is exaggerated upon MIA induction. Expansion of GFP+ drHSCs one day post MIA is therefore likely responsible for subsequent expansion of Flk2+ MPP4s one day later, and sustained expansion of both populations was also observed postnatally. Our analysis of the initial response of HSPCs to interferons using the Mx1-Cre model similarly confirms that HSCs drive the initial response to pIC-induced prenatal inflammation. Together, our data indicate that inflammation during the fetal period invokes a lymphoid-biased response *in situ* by activating lymphoid-biased HSCs that subsequently drive the downstream expansion of lymphoid-biased progenitors.

In contrast to the lymphoid-biased hematopoietic response observed *in situ*, adoptive transfer experiments revealed that Tom+ HSCs exhibited higher GM chimerism upon transplantation, suggesting the possibility that MIA also activated Tom+ HSCs and induced myeloid bias. However, neither expansion of Tom+ HSCs nor myeloid bias was observed *in situ.* Similarly, whereas enhanced lymphoid differentiation was observed ex vivo and *in situ* in response to MIA, MIA did not exaggerate lymphoid bias in GFP+ drHSCs upon transplantation. This discrepancy in the *in situ* response as compared to the output of HSCs upon transplantation could be due to exposure to different microenvironmental cues in an irradiated, adult environment (Guo et al., 2017). Alternatively, these data suggest that prenatal inflammation does not change the cell-intrinsic bias of fetal HSCs, but instead functions to activate and expand specific subsets with the fetal liver microenvironment. Interestingly, serial transplantation resulted in rapid exhaustion of Tom+ HSCs, but not GFP+ drHSCs. This observation is consistent with prenatal inflammation inducing abnormal persistence of the GFP+ drHSC, potentially at the expense of persistence of some fraction of the Tom+ HSCs. Fundamental differences in the composition of adult and fetal hematopoietic compartments (Beaudin *et al*., 2016; Benz et al., 2012; Böiers et al., 2013; Crisan et al., 2016; Li et al., 2020), may underlie divergent hematopoietic trajectories between fetal and adult hematopoiesis in response to inflammation. Specifically, activation of a transient, lymphoid-biased fetal HSC may reflect the increased sensitivity of transient fetal progenitors to prenatal inflammation as a mechanism to prime the developing immune system. This hypothesis is consistent with the concept of layered immunity (Herzenberg, 1989), whereby transient precursors give rise to specialized subsets of immune cells, such as B1 B-cells, that may be more receptive to insult during the prenatal period. Susceptibility of adult HSC precursors, and the long-term effects of prenatal inflammation on their self-renewal capability, output, and response to infection, requires additional investigation.

Our single-cell analysis of over 23,000 fetal single HSPCs confirmed the expansion of both HSC-like progenitors and lymphoid-primed progenitors in response to prenatal inflammation. At least two clusters, Cluster (C) 2 and C9, resembled HSCs in both transcriptional profile and proliferative state, but only one of these populations, C9, overlapped transcriptionally with the GFP+ drHSC and was expanded in response to MIA. C9 also stood out as the only cluster to exhibit a strong cytokine-responsive transcriptional profile, and increased inflammatory gene expression in response to MIA. Cytokine analysis revealed upregulation of IFN-α in the FL, and C9 exhibited a robust transcriptional response to interferon. Surprisingly, upregulation of type I interferons did not increase non-specific Sca1 expression in the fetus, as observed in adult hematopoiesis (King and Goodell, 2011). In contrast, C2, which overlaps transcriptionally with the Tom+ HSC, exhibited a more “adult-like” HSC profile, as revealed by expression of more canonical HSC genes including *Mllt3* and *Meis1*, among others. Furthermore, C2 contracted in response to MIA, and expressed little to no inflammatory gene signature. Our transcriptional analysis mirrors our *in vivo* data, whereby we identified an HSC-like cluster, overlapping with the GFP+ drHSC, that both expands and exhibits a robust transcriptional response to cytokines that are upregulated within the fetal microenvironment in response to prenatal inflammation.

In addition to an HSC-like cluster, our single-cell transcriptional profiling also identified a highly lymphoid-biased cluster, C10, that overlapped transcriptionally with the GFP+ drHSC signature. Not only did MIA cause significant expansion of C10, but we also observed upregulated expression of lymphoid-associated genes such as *Ighm, Chchd10, Mzb1,* and *Vpreb 1/3* in response to MIA. Notably, many of these genes are associated with B-cell and innate-like B-cell lineage commitment, possibly skewing towards production of hyperactivated innate-like B-cells in the postnatal period. Whether increased expression of lymphoid-associated genes in fetal progenitors ultimately contributes to a hyperactivated phenotype that was observed in lymphocyte progeny postnatally, remains to be determined. Nonetheless, this profile parallels our *in vivo* observation of specific expansion not only of the GFP+ drHSC, but also of lymphoid-biased progenitors in response to MIA, and provides additional evidence that prenatal inflammation drives lymphoid bias during the fetal and postnatal period.

The MIA model was used previously to examine how prenatal inflammation affects brain development during a “critical window” of synapse formation (Meyer *et al*., 2009). A critical window is defined as a developmental window during which extrinsic or intrinsic inputs can shape the phenotype of the adult organism. Here, we use the same approach to establish a framework for a critical window for hematopoietic and immune development wherein extrinsic input, in the form of *in utero* inflammation, shapes the establishment and output of the hematopoietic and immune systems across ontogeny. Although it was previously reported that under homeostatic conditions the GFP+ drHSC fails to persist into adulthood (Beaudin *et al*., 2016; Beaudin and Forsberg, 2016), here we observed the persistence of some phenotypic GFP+ drHSCs into adulthood under homeostatic conditions. We believe this is due to increased basal inflammation within our mouse colony, which fits with our observation of persistence of GFP+ drHSCs induced by inflammation with MIA, as well as basal labeling by Mx1-Cre in the absence of MIA. Nonetheless, heightened inflammation induced by MIA during the fetal period drove increased proliferation, expansion, and enhanced persistence of GFP+ drHSCs postnatally and into adulthood. Concomitant with postnatal expansion of a lymphoid-biased GFP+ drHSC, we observed the specific expansion and hyperactivation of fetal-derived innate-like lymphocyte compartments; the innate-like B1 B-cell compartment was not only increased in cell number, but produced more natural antibody and cytokine on a per-cell basis. Fetal B-cells did not expand in direct response to prenatal inflammation induced by MIA, nor was their response to interferons detectable using the Mx1-Cre model as a fate-mapping approach, indicating that their expansion postnatally was not due to a direct response to prenatal inflammation. Although we cannot directly track progeny of the GFP+ drHSC *in situ*, due to limitations of the FlkSwitch model (Boyer *et al*., 2012), the coincidence of expanded and activated GFP+ drHSCs, and their innate-like lymphocyte progeny in response to prenatal inflammation strongly suggests a causal relationship. These data suggest that prenatal inflammation during fetal development can shape postnatal hematopoietic and immune output. We did not directly test the effect of an expanded and hyperactivated B1 B-cell compartment on offspring immunity. Expanded and hyperactivated B1 B-cells and increased natural antibody couldx be expected to protect neonates from severe infections such as sepsis (Baumgarth, 2021; Smith and Baumgarth, 2019), although elevated IL-10 levels might also increase susceptibility (Romagnoli et al., 2001; Ye et al., 2017). Together, our results illuminate how *in utero* inflammation can influence immune trajectories by reshaping hematopoiesis during development. Our discovery has significant implications for understanding the ontogeny of many immune-related disorders with poorly understood etiology, including diseases of tolerance, autoimmunity, and clonal hematopoiesis. Future studies should directly address how lasting changes to immune output and function modulate susceptibility to infection and disease.

## Acknowledgements

We thank Dr. T. Boehm for the use of the Flt3-Cre mice, David Gravano and the UC Merced Stem Cell Instrumentation Foundry, James Marvin and the University of Utah Flow Cytometry Core, Bari Nazario for flow cytometry support, and Jasmine Posada for technical assistance. The sequencing was carried out by the DNA Technologies and Expression Analysis Core at the UC Davis Genome Center, supported by NIH Shared Instrumentation Grant 1S10OD010786-01. This work was supported by an NIH/NHBLI award K01HL130753 to AEB, the Pew Biomedical Scholars award to AEB, and the Hellman Fellows Award to AEB; NIH/NIDDK R01DK100917 to ECF; and Max Planck Society and the ERC-Stg-2017 (VitASTEM) Research to NC-W. This work was supported by CIRM facilities awards CL1-00506 and FA1-00617 to UCSC and the DoD Research and Education Program for HBCU/MSI Instrumentation Grant W911NF1910529, as well as the National Center for Research Resources of the NIH under award number 1S10RR026802-01. DAL is supported by NIH/NICHD training grant T32HD007491. GEH is supported by Howard Hughes Medical Institute Gilliam Fellowship (GT11560). AEB is the recipient of an NIH/NHLBI R01HL147081.

## Author contributions

AEB was responsible for study conceptualization, experimental design, data collection, data analysis, data visualization, preparation of the manuscript, and supervision of the work. ECF was responsible for study conceptualization. DAL and ACA were responsible for experimental design, data collection, data analysis, data visualization, and preparation of the manuscript. EJL, CDV and GEH performed experiments and analyzed data. MR-M and PP performed computational analysis and assisted with data visualization. NC-W performed computational analysis, assisted with data visualization and manuscript preparation. AEB provided funding for this work.

## STAR METHODS

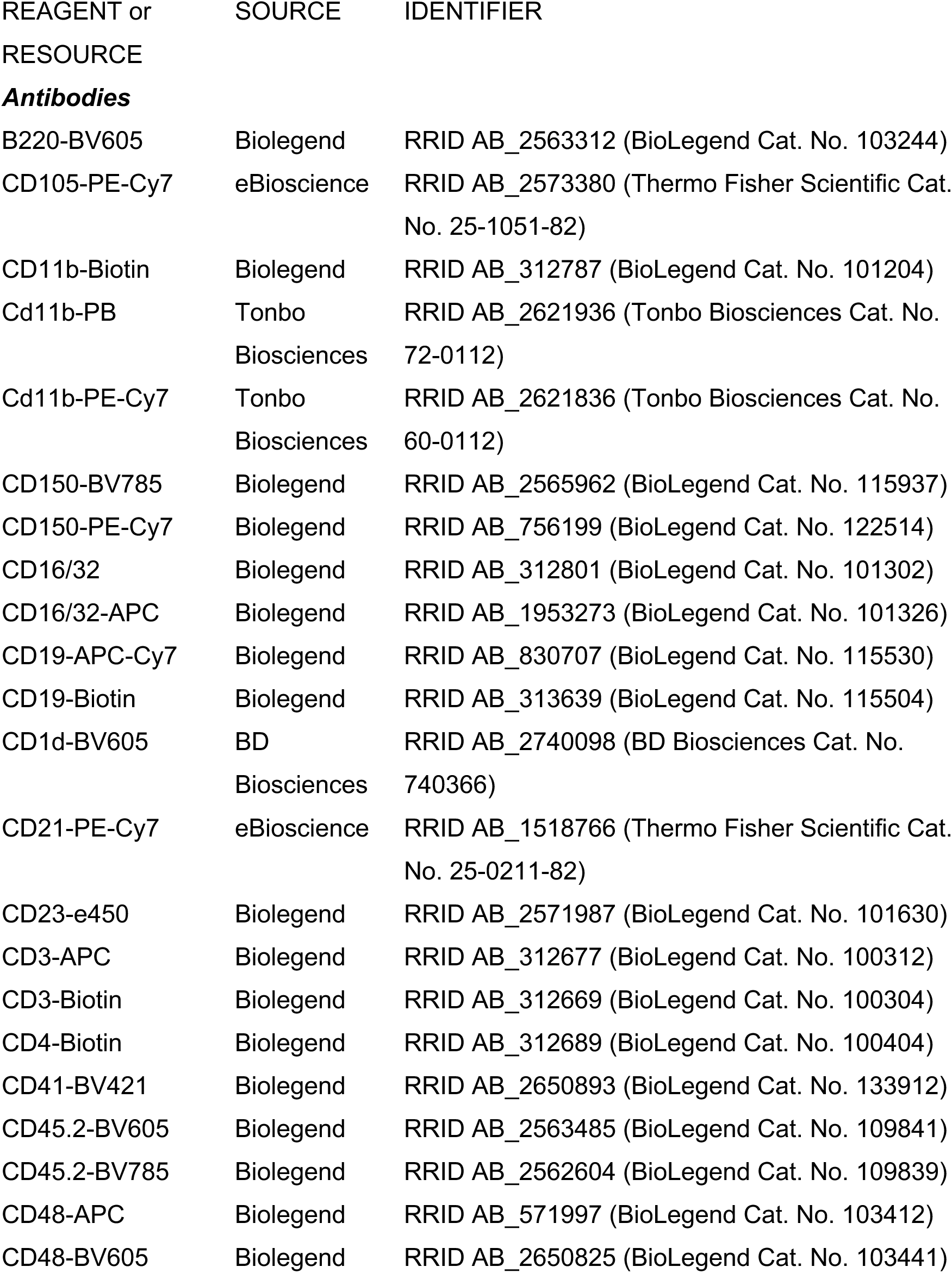
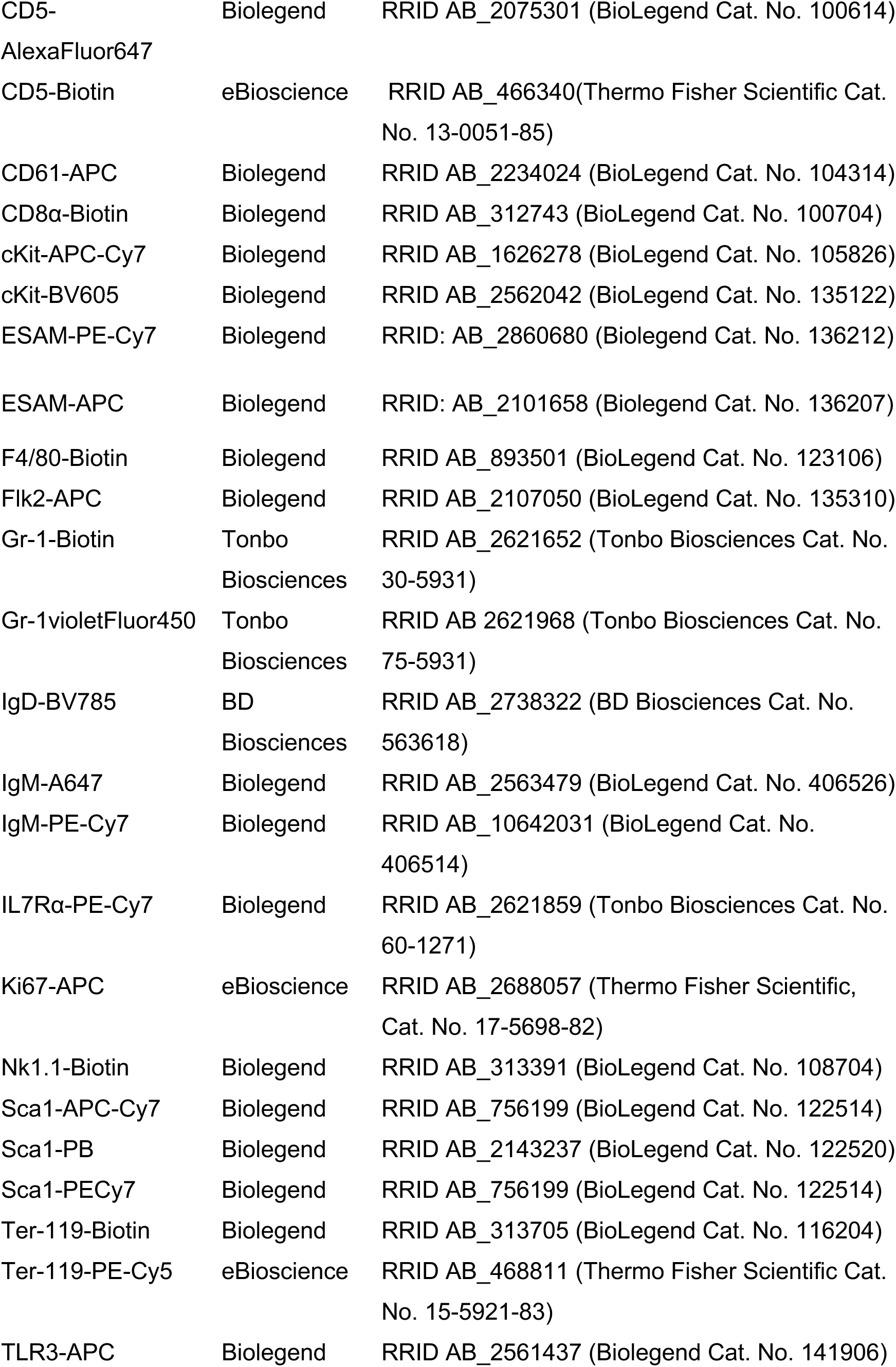
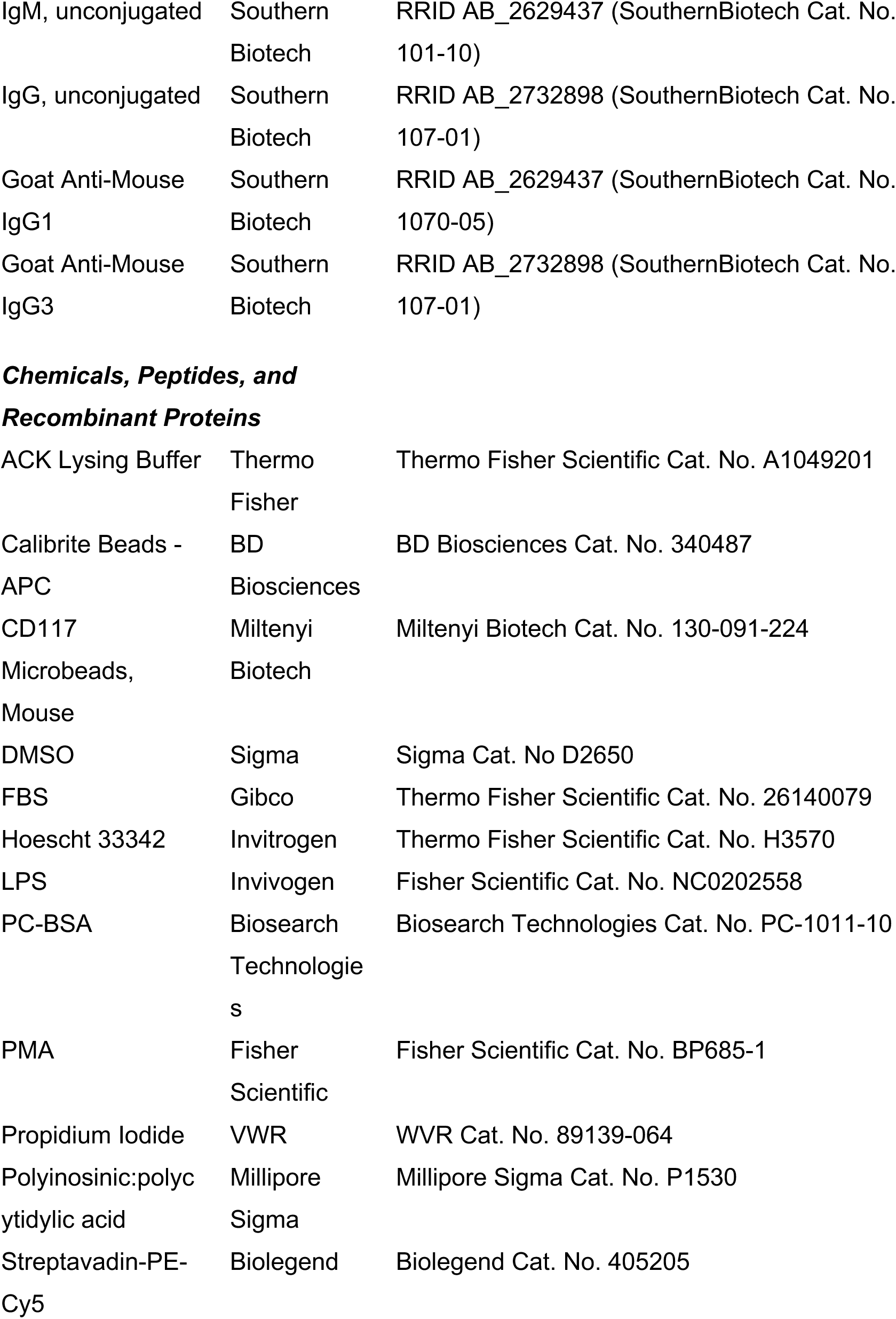
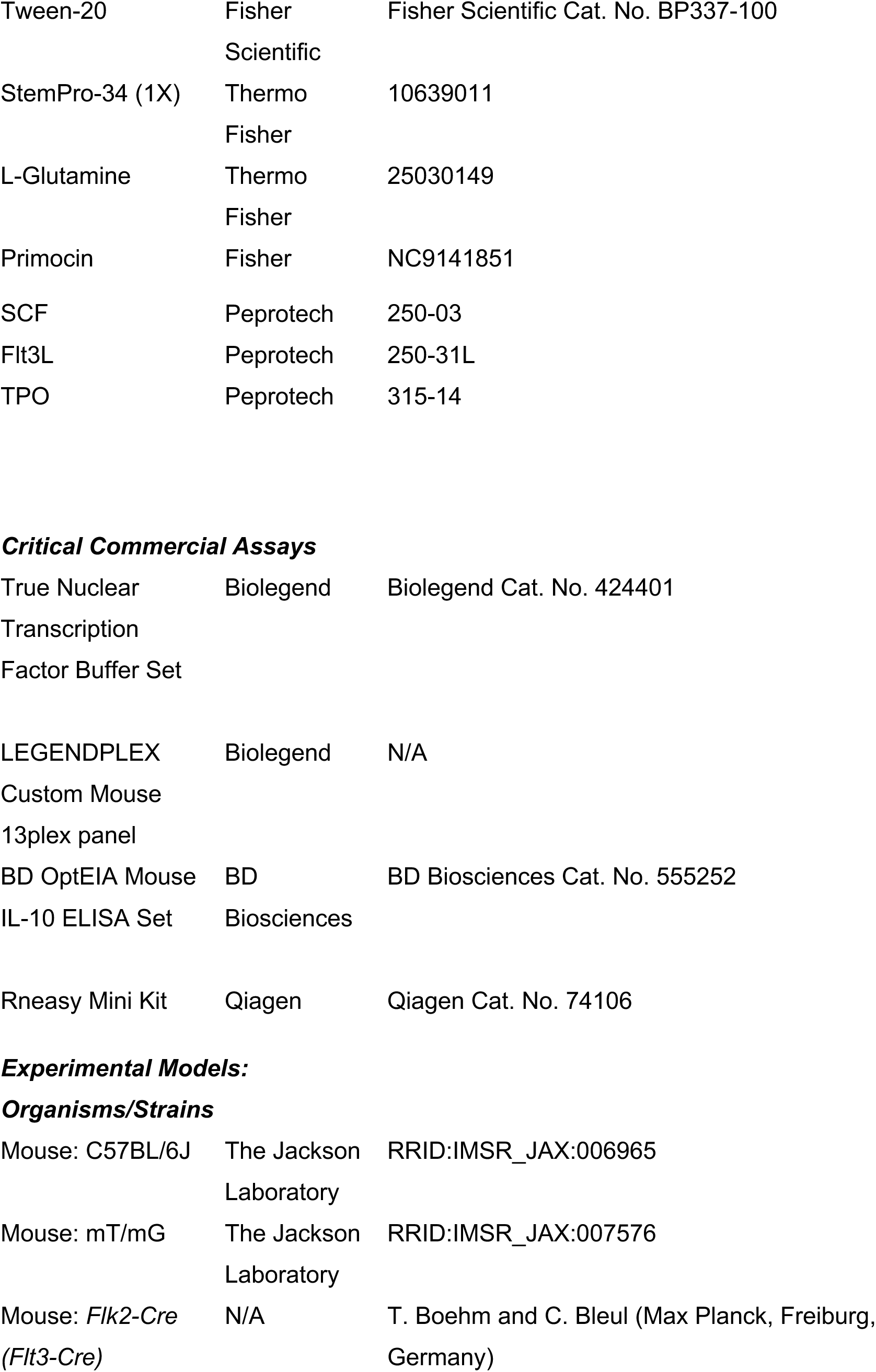
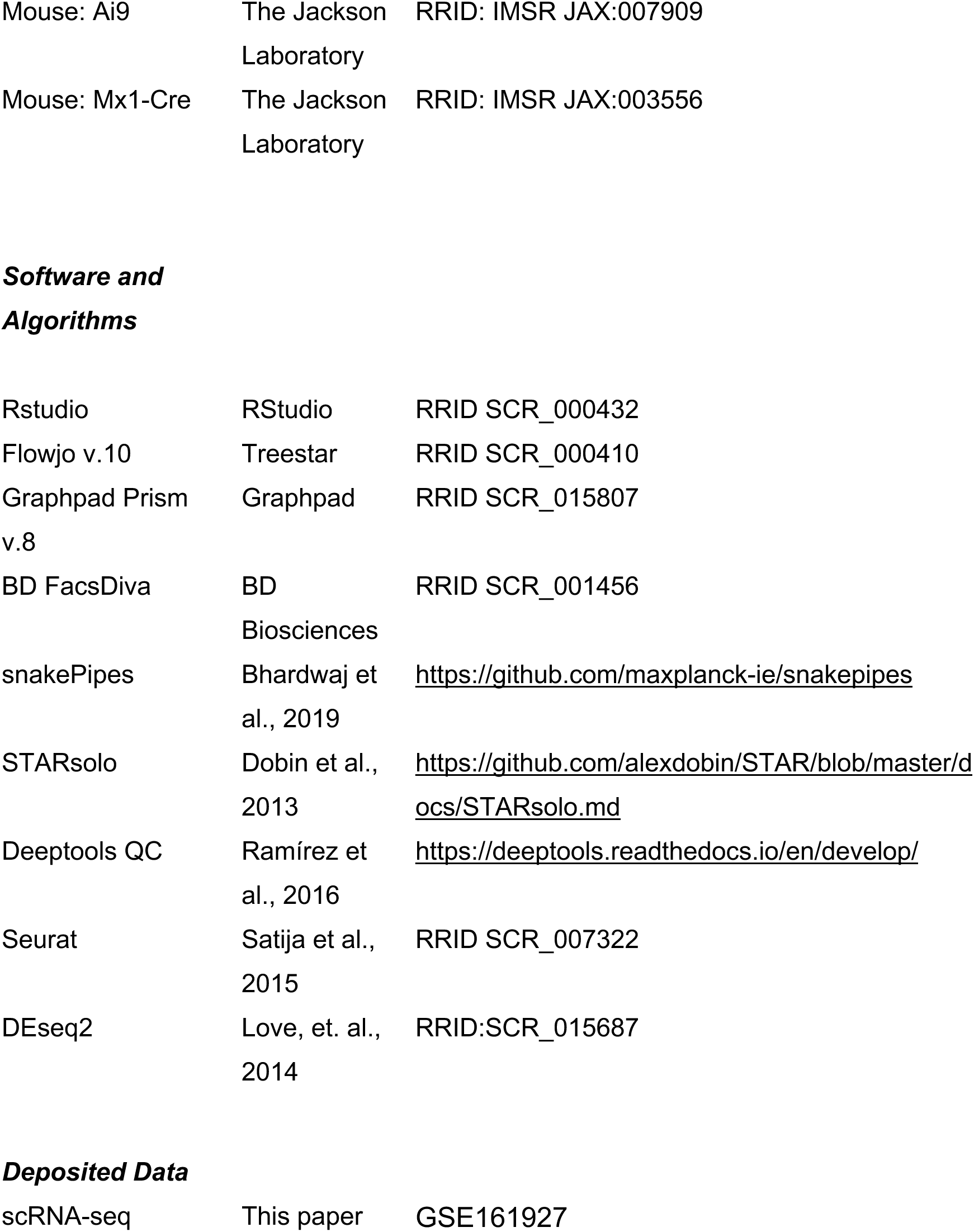

| REAGENT or RESOURCE | SOURCE | IDENTIFIER |
| --- | --- | --- |
| ***Antibodies*** |  |  |
| B220-BV605 | Biolegend | RRID AB_2563312 (BioLegend Cat. No. 103244) |
| CD105-PE-Cy7 | eBioscience | RRID AB_2573380 (Thermo Fisher Scientific Cat. No. 25-1051-82) |
| CD11b-Biotin | Biolegend | RRID AB_312787 (BioLegend Cat. No. 101204) |
| Cd11b-PB | Tonbo Biosciences | RRID AB_2621936 (Tonbo Biosciences Cat. No. 72-0112) |
| Cd11b-PE-Cy7 | Tonbo Biosciences | RRID AB_2621836 (Tonbo Biosciences Cat. No. 60-0112) |
| CD150-BV785 | Biolegend | RRID AB_2565962 (BioLegend Cat. No. 115937) |
| CD150-PE-Cy7 | Biolegend | RRID AB_756199 (BioLegend Cat. No. 122514) |
| CD16/32 | Biolegend | RRID AB_312801 (BioLegend Cat. No. 101302) |
| CD16/32-APC | Biolegend | RRID AB_1953273 (BioLegend Cat. No. 101326) |
| CD19-APC-Cy7 | Biolegend | RRID AB_830707 (BioLegend Cat. No. 115530) |
| CD19-Biotin | Biolegend | RRID AB_313639 (BioLegend Cat. No. 115504) |
| CD1d-BV605 | BD Biosciences | RRID AB_2740098 (BD Biosciences Cat. No. 740366) |
| CD21-PE-Cy7 | eBioscience | RRID AB_1518766 (Thermo Fisher Scientific Cat. No. 25-0211-82) |
| CD23-e450 | Biolegend | RRID AB_2571987 (BioLegend Cat. No. 101630) |
| CD3-APC | Biolegend | RRID AB_312677 (BioLegend Cat. No. 100312) |
| CD3-Biotin | Biolegend | RRID AB_312669 (BioLegend Cat. No. 100304) |
| CD4-Biotin | Biolegend | RRID AB_312689 (BioLegend Cat. No. 100404) |
| CD41-BV421 | Biolegend | RRID AB_2650893 (BioLegend Cat. No. 133912) |
| CD45.2-BV605 | Biolegend | RRID AB_2563485 (BioLegend Cat. No. 109841) |
| CD45.2-BV785 | Biolegend | RRID AB_2562604 (BioLegend Cat. No. 109839) |
| CD48-APC | Biolegend | RRID AB_571997 (BioLegend Cat. No. 103412) |
| CD48-BV605 | Biolegend | RRID AB_2650825 (BioLegend Cat. No. 103441) |
| CD5-AlexaFluor647 | Biolegend | RRID AB_2075301 (BioLegend Cat. No. 100614) |
| CD5-Biotin | eBioscience | RRID AB_466340(Thermo Fisher Scientific Cat. No. 13-0051-85) |
| CD61-APC | Biolegend | RRID AB_2234024 (BioLegend Cat. No. 104314) |
| CD8α-Biotin | Biolegend | RRID AB_312743 (BioLegend Cat. No. 100704) |
| cKit-APC-Cy7 | Biolegend | RRID AB_1626278 (BioLegend Cat. No. 105826) |
| cKit-BV605 | Biolegend | RRID AB_2562042 (BioLegend Cat. No. 135122) |
| ESAM-PE-Cy7 | Biolegend | RRID: AB_2860680 (Biolegend Cat. No. 136212) |
| ESAM-APC | Biolegend | RRID: AB_2101658 (Biolegend Cat. No. 136207) |
| F4/80-Biotin | Biolegend | RRID AB_893501 (BioLegend Cat. No. 123106) |
| Flk2-APC | Biolegend | RRID AB_2107050 (BioLegend Cat. No. 135310) |
| Gr-1-Biotin | Tonbo Biosciences | RRID AB_2621652 (Tonbo Biosciences Cat. No. 30-5931) |
| Gr-1violetFluor450 | Tonbo Biosciences | RRID AB 2621968 (Tonbo Biosciences Cat. No. 75-5931) |
| IgD-BV785 | BD Biosciences | RRID AB_2738322 (BD Biosciences Cat. No. 563618) |
| IgM-A647 | Biolegend | RRID AB_2563479 (BioLegend Cat. No. 406526) |
| IgM-PE-Cy7 | Biolegend | RRID AB_10642031 (BioLegend Cat. No. 406514) |
| IL7Rα-PE-Cy7 | Biolegend | RRID AB_2621859 (Tonbo Biosciences Cat. No. 60-1271) |
| Ki67-APC | eBioscience | RRID AB_2688057 (Thermo Fisher Scientific, Cat. No. 17-5698-82) |
| Nk1.1-Biotin | Biolegend | RRID AB_313391 (BioLegend Cat. No. 108704) |
| Sca1-APC-Cy7 | Biolegend | RRID AB_756199 (BioLegend Cat. No. 122514) |
| Sca1-PB | Biolegend | RRID AB_2143237 (BioLegend Cat. No. 122520) |
| Sca1-PECy7 | Biolegend | RRID AB_756199 (BioLegend Cat. No. 122514) |
| Ter-119-Biotin | Biolegend | RRID AB_313705 (BioLegend Cat. No. 116204) |
| Ter-119-PE-Cy5 | eBioscience | RRID AB_468811 (Thermo Fisher Scientific Cat. No. 15-5921-83) |
| TLR3-APC | Biolegend | RRID AB_2561437 (Biolegend Cat. No. 141906) |
| IgM, unconjugated | Southern Biotech | RRID AB_2629437 (SouthernBiotech Cat. No. 101-10) |
| IgG, unconjugated | Southern Biotech | RRID AB_2732898 (SouthernBiotech Cat. No. 107-01) |
| Goat Anti-Mouse IgG1 | Southern Biotech | RRID AB_2629437 (SouthernBiotech Cat. No. 1070-05) |
| Goat Anti-Mouse IgG3 | Southern Biotech | RRID AB_2732898 (SouthernBiotech Cat. No. 107-01) |
| ***Chemicals, Peptides, and Recombinant Proteins*** |  |  |
| ACK Lysing Buffer | Thermo Fisher | Thermo Fisher Scientific Cat. No. A1049201 |
| Calibrite Beads - APC | BD Biosciences | BD Biosciences Cat. No. 340487 |
| CD117 Microbeads, Mouse | Miltenyi Biotech | Miltenyi Biotech Cat. No. 130-091-224 |
| DMSO | Sigma | Sigma Cat. No D2650 |
| FBS | Gibco | Thermo Fisher Scientific Cat. No. 26140079 |
| Hoescht 33342 | Invitrogen | Thermo Fisher Scientific Cat. No. H3570 |
| LPS | Invivogen | Fisher Scientific Cat. No. NC0202558 |
| PC-BSA | Biosearch Technologies | Biosearch Technologies Cat. No. PC-1011-10 |
| PMA | Fisher Scientific | Fisher Scientific Cat. No. BP685-1 |
| Propidium Iodide | VWR | WVR Cat. No. 89139-064 |
| Polyinosinic:polycytidylic acid | Millipore Sigma | Millipore Sigma Cat. No. P1530 |
| Streptavidin-PE-Cy5 | Biolegend | Biolegend Cat. No. 405205 |
| Tween-20 | Fisher Scientific | Fisher Scientific Cat. No. BP337-100 |
| StemPro-34 (1X) | Thermo Fisher | 10639011 |
| L-Glutamine | Thermo Fisher | 25030149 |
| Primocin | Fisher | NC9141851 |
| SCF | Peptotech | 250-03 |
| Flt3L | Peptotech | 250-31L |
| TPO | Peptotech | 315-14 |
| ***Critical Commercial Assays*** |  |  |
| True Nuclear Transcription Factor Buffer Set | Biolegend | Biolegend Cat. No. 424401 |
| LEGENDPLEX Custom Mouse 13plex panel | Biolegend | N/A |
| BD OptEIA Mouse IL-10 ELISA Set | BD Biosciences | BD Biosciences Cat. No. 555252 |
| Rneasy Mini Kit | Qiagen | Qiagen Cat. No. 74106 |
| ***Organisms/Strains*** |  |  |
| Mouse: C57BL/6J | The Jackson Laboratory | RRID:IMSR_JAX:006965 |
| Mouse: mT/mG | The Jackson Laboratory | RRID:IMSR_JAX:007576 |
| Mouse: *Flk2-Cre* (*Flt3-Cre*) | N/A | T. Boehm and C. Bleul (Max Planck, Freiburg, Germany) |
| Mouse: Ai9 | The Jackson Laboratory | RRID: IMSR JAX:007909 |
| Mouse: Mx1-Cre | The Jackson Laboratory | RRID: IMSR JAX:003556 |
| ***Software and Algorithms*** |  |  |
| Rstudio | RStudio | RRID SCR_000432 |
| Flowjo v.10 | Treestar | RRID SCR_000410 |
| Graphpad Prism v.8 | Graphpad | RRID SCR_015807 |
| BD FacsDiva | BD Biosciences | RRID SCR_001456 |
| snakePipes | Bhardwaj et al., 2019 | https://github.com/maxplanck-ie/snakepipes |
| STARsolo | Dobin et al., 2013 | https://github.com/alexdobin/STAR/blob/master/docs/STARsolo.md |
| Deeptools QC | Ramírez et al., 2016 | https://deeptools.readthedocs.io/en/develop/ |
| Seurat | Satija et al., 2015 | RRID SCR_007322 |
| DEseq2 | Love, et. al., 2014 | RRID:SCR_015687 |
| ***Deposited Data*** |  |  |
| scRNA-seq | This paper | GSE161927 |

## RESOURCE AVAILABILITY

### Lead Contact

- Further information and requests for resources and reagents should be directed to and will be fulfilled by the Lead Contact, Anna E. Beaudin

### Materials Availability

- This study did not generate new unique reagents.

### Data and Code Availability

#### Accession numbers

The accession number for GEO where the RNA-seq datasets generated in this study are available is GSE161927.

## EXPERIMENTAL MODEL AND SUBJECT DETAILS

### Mouse models and husbandry

8-12-week-old female C57BL/6 (RRID: IMSR_JAX:000664) were mated to male Flkswitch mice (Boyer et al., 2011, 2012; Epelman et al., 2014; Hashimoto et al., 2013, Beaudin, et al., 2016). Flkswitch mice were generated by crossing Flk2-Cre mice (Benz et al., 2008) to mTmG mice (Muzumdar et al., 2007) and gestation day was confirmed by the presence of a sperm plug. Flkswitch males were used because the Flk-Cre transgene is only transmitted through the Y chromosome. At gestation day 14.5, pregnant dams were weighed and injected with 20mg/kg Polyinosinic:polycytidylic acid (pIC). For fetal timepoints, pregnant dams were euthanized, and fetuses were dissected from the uterine horn. Fetal liver GFP+ (Flkswitch) expression was confirmed by microscopy in fetuses or by peripheral blood analysis at postnatal timepoints, Postnatal day (P)14 and 8-week-old adult. For transplantation assays, FlkSwitch mice were used as donors for cell isolation and 8- to 12-week-old WT C57BL/6 were used as recipients. Sex of recipients was random and split evenly between male and female. All mice were maintained in the University of California, Merced vivarium according to Institutional Animal Care and Use Committee (IACUC)-approved protocols.

### Cell isolation and identification by flow cytometry

Fetal livers were dissected and pipetted gently in staining media to form a single cell suspension. Adult and P14 BM cells isolated by dissecting long bones and using a mortar and pestle to gently crush bones in “staining media” (1XPBS, 2%FBS, .5mM EDTA) and bone marrow was extracted by gently pipetting. Cells were filtered through a 70uM filter. Cell populations were analyzed using a four-laser FACSAria III (BD Biosciences). Cells were sorted on The FACSAriaII (BD Biosciences) or III. All flow cytometric analysis was done using FlowJo™. Hematopoietic and mature blood cell populations were identified as follows: Lineage dump or “Lin” for all fetal liver populations (CD3, CD4, CD5, CD8, CD19, Ter-119, Nk1.1, Gr-1, F4/80), Lin for adult bone marrow populations (CD3, CD4, CD5, CD8, Cd11b, CD19, Ter-119, Nk1.1, Gr-1, F4/80); <u>LT-HSCs</u> (Lin-, CD45+, cKit+, Sca1+, Flk2-, CD48-, CD150+), <u>ST-HSCs</u> (Lin-, CD45+, cKit+, Sca1+, Flk2-, CD48-, CD150-), <u>Tom+ aHSCs</u>, (Lin-, CD45+, cKit+, Sca1+, CD150+, Tom+), <u>GFP+ drHSCs</u> (Lin-, CD45+, cKit+, Sca1+, CD150+, GFP+), <u>MPP2</u> (Lin-, CD45+, cKit+, Sca1+, Flk2-, CD48+, CD150+), <u>MPP3</u> (Lin-, CD45+, cKit+, Sca1+, Flk2-, CD48+ CD150-), <u>MPP4</u> (Lin-, CD45+, CKit+, Sca1+, Flk2+, CD48+, CD150-), Granulocyte-Macrophage Progenitor “<u>GM</u>” (Lin-CKit+, CD41-, CD16/32+), Megakaryocyte Progenitor “<u>MP</u>”, Erythrocyte progenitor “<u>EP</u>”; Peritoneal Cavity: <u>B1a</u> (CD3-, F4/80-Gr1-,Ter119-, CD19+ CD5+), <u>B1b</u> (CD3-, F4/80-Gr1-,Ter119-, CD19+ CD5-, CD11b+), <u>B2</u> (CD3-, F4/80-Gr1-,Ter119-, CD19+ CD5-, CD11b-); Splenic B cells: Marginal Zone B-cells “<u>MZ</u>” (CD3-, F4/80-Gr1-,Ter119-, IgM+, CD19+, CD21+ CD23-), Follicular Zone “<u>FZ</u>” B-cells (CD3-, F4/80-Gr1-,Ter119-, IgM+, CD19+, CD21- CD23+).

### Proliferation of HSCs and MPPs

Fetal liver cells were processed into a single cell suspension and cKit-enriched using CD117 MicroBeads (Miltenyi Biotec, San Diego, CA, USA). The cKit-enriched population was stained with an antibody cocktail for surface markers of hematopoietic stem and progenitor cells. Cells were then fixed and permeabilized with the True-Nuclear Transcription buffer set (Biolegend) and then stained with Ki67-APC (Invitrogen, Carlsbad, CA, USA) and Hoescht 33342 (Invitrogen).

### Culture of Tom+ HSC or GFP+ drHSC to assess HSPC generation

E14.5 FL GFP+ drHSCs and Tom+ HSCs were sorted form FlkSwitch embryos 2 hours post *in vivo* saline or MIA treatment. 450 Tom+ HSCs or GFP+ drHSCs were cultured at 37°C with 5% CO_2_ in StemPro^®^-34 SFM supplemented with 2mM L-Glutamine, 50 ng/mL SCF, 30 ng/mL Flt3L, 25 ng/mLTPO, and 100 ug/mL Primocin. Cells were harvested after 24 hours, stained for HSPC surface cell markers, and analyzed using an Aurora Spectral Analyzer (Cytek). MPP3 and MPP4 populations were defined as Lin-cKit+Sca1+CD48+CD150-ESAM+ and cKit+Sca1+CD48+CD150-ESAM-, respectively (Beaudin et al., 2014).

### Transplantation assays

Transplantations were performed by is sorting GFP+ drHSCs and Tom+ HSCs from fetal liver. Recipient C57BL/6 mice (8-12 weeks) were lethally irradiated using 1000 cGy (split dose, Precision X-Rad 320). 5×10^6^ whole bone marrow cells from untreated age matched C57BL/6 and 200 sorted GFP+ drHSCs or Tom+ HSCs wells were diluted in PBS and transplanted via retroorbital injection using a 1mL tuberculin syringe in a volume of 100-200 uL. Peripheral blood chimerism was determined in recipients by blood collection via cheek bleeds every 4 weeks for 16 weeks and cells were analyzed by flow cytometry using the LSRII (BD Biosciences). Long-term multilineage reconstitution (LTMR) was defined as chimerism >1% in all mature blood lineages. At 18 weeks, recipients were euthanized, and BM populations were assessed for chimerism by flow cytometry.

### 10x Chromium single-cell sequencing of HSPCs

Pregnant dams from C57BL/6 Flkswitch crosses were exposed to pIC or saline at E14.5. E15.5 fetal liver cells were pooled, cKit-enriched, and sorted for HSPCs (CD45+, Lin-, cKit+ Sca1+). In order to get adequate cell numbers for analysis, at least 9 litters/condition (with 3-8 fetal livers/litter) were used for sorts. Sorted HSPCs were frozen at -80°C in FBS and 15% DMSO in cryovials until all samples were collected. Frozen cells were thawed in a 37°C water bath for 2-3 minutes and washed and resuspended in 10% FBS and assessed for 60-70% viability before processing. A sample library was generated using Chromium 10x V3 and run using NovaSeq.

### scRNA-seq data pre-processing and quality control

Raw UMI-based data files were mapped against the mm10 reference genome using the scRNAseq tool from the bioinformatics pipeline snakePipes, with the 10xV3 mode (Bhardwaj et al., 2019). In this tool, STARsolo was used to i) map, ii) UMI-deduplicate and iii) count reads, to create the BAM files and a Seurat object with the gene counts (Dobin et al., 2013). The quality of this data was checked by running Deeptools QC (Ramirez et al., 2016). After the preprocessing of data, R package Seurat was used to perform the single-cell RNA-seq analysis (Satija et al., 2015). The Seurat object was imported, and a cell filtering was performed, selecting the cells which i) contained more than 75,000 counts and ii) those that expressed above 2,000 genes to avoid empty droplets. Moreover, low-quality and dying cells with a percentage of mitochondrial mRNA higher than 20% were filtered out.

### scRNA-seq data clustering and gene expression

A log-normalization and scaling of the data was applied prior to the linear dimension reduction with the Principal Component Analysis (PCA). After applying the JackStraw and Elbow plot procedures from Seurat, a clustering analysis was performed selecting the first 20 PCs, resulting in cell communities that were visualized with the Uniform Manifold Approximation and Projection (UMAP) technique (Becht et al., 2018). From those, three clusters were filtered out based on the doublets scoring calculated with the doubletCells function from scran package (Lun et al., 2016), the enrichment of LSK and LS-K signatures described previously (Klimmeck et al., 2014) applying the AddModuleScore function and the presence of differentiated cells markers, detected with the FindAllMarkers function (min.pct = 0, logfc.threshold = 0.15). A total of 23,505 cells distributed in 14 clusters were further analyzed, corresponding 11,977 and 11,528 cells to the pIC and Saline conditions, respectively. A heatmap with the expression of the top seven differentially expressed genes per cluster was performed with DoHeatmap function. The percentage of cells in each condition per cluster was visualized in a barplot, assessing the significance with Fisher tests. Cell cycle phase alignment was performed by calculating the G1 and G2M scores defined by the cyclone function from scran package, using default parameters (Scialdone et al., 2015). The enrichment of the eGFP, tdTomato signatures generated using reference data set from E14.5 fetal liver (Beaudin et al., 2016) and the also generated HSC and MPP gene expression signatures (Cabezas-Wallscheid et al., 2014) were depicted by the AddModuleScore function. The resulting enrichment was represented in the UMAP projection with the FeaturePlot function and in violin plots, calculating the significance per cluster with respect to all scores using a Wilcoxon signed rank test with one-sided alternative hypothesis. In the same way, the enrichment of the reactome database (reactome.org) pathways integral to cytokine, interferon and interleukin signaling in the immune system were represented individually in the single cell data as violin plots. Generated lists of cluster markers by the FindMarkers function (min.pct=0.25, logfc.threshold=0.25) were used for gene set enrichment analysis (GSEA) in the already published ranked set of differentially expressed genes between Tom+ HSC and GFP+ drHSCs (Beaudin et al., 2016), using the GSEA and gseaplot2 functions (Yu, 2021).

### TLR3 intracellular and extracellular staining

8-12-week-old female C57BL/6 female mice were mated to C57BL/6 males and treated with 20mg/kg pIC or saline at E14.5. Females were sacrificed 2hrs later and FLs were harvested from embryos and processed as described above and. Cells were then stained using an HSPC antibody cocktail before being fixed and permeabilized with the True-Nuclear Transcription buffer set (Biolegend) and stained with TLR3-APC (Biolegend). Flow cytometric analysis was performed using an Aurora Spectral Analyzer (Cytek).

### Inducible Mx-1 Cre lineage tracing

8-12-week-old female Ai9 reporter mice (RRID: IMSR JAX:007909) were mated to male Mx1-Cre mice (RRID: IMSR JAX:003556) and treated with 20mg/kg pIC or saline (control) at E14.5. After 1hr, 24 hrs, or 72 hrs post-MIA, FLs were processed as described above and cells were stained with antibody cocktails for HSPC, MkP (Lin-CD45+cKit+Sca1-CD150+CD41+), GMP (Lin-CD45+cKit+Sca1-CD150-FcγRIII/II+), Pre-GM (Lin-CD45+cKit+Sca1-CD150-FcγRIII/II-CD105-), Pre-MegE (Lin-CD45+cKit+Sca1-CD150+FcγRIII/II-CD105^lo/-^), Pre CFU-E (Lin-CD45+cKit+Sca1-CD150+FcγRIII/II-CD105+), as defined by (Pronk et al., 2007), and B1a, and B1b populations. Flow cytometric analysis was performed using an Aurora Spectral Analyzer (Cytek). Frequency of label (%TdTomato+) was determined by taking the average background labeling in saline-treated controls and subtracting it from the frequency of label for each population in pIC-treated mice.

### Quantitation of cytokines in fetal amniotic fluid and liver

Fetal amniotic fluid was collected from amniotic sacs using a 1 mL tuberculin syringe. Fetuses were dissected and individual fetal livers were homogenized and pelleted to collect supernatant. Supernatant was frozen at -80°C and samples were analyzed on an LSRII (BD Biosciences) following manufacturer recommendations using a custom LEGENDplex^TM^ (Biolgegend) bead-based immunoassay for the following cytokines: IL-23, IL-1α, IFNγ, TNF-α, IL-6, IL-1β, IL-10, IL-17A, IL-27, IFNβ, IFNα, and GM-CSF. Data were analyzed with LEGENDplex^TM^ online analysis platform. https://legendplex.qognit.com/.

### Isolation and stimulation of postnatal day 21 peritoneal B cells for IL-10 quantitation

Peritoneal washouts were performed by injecting 1mL of cold media + 2% FBS into the peritoneal cavity (PEC) of P21 pups. Mice were shaken 10-15 times and media was pipetted out of the abdomen via a cut beneath the sternum. Multiple washes were performed (for a total of 3 washes) by adding more media through cut under the sternum. Samples were filtered and pooled together for sorting. Cell populations were sorted and resuspended in sterile 1X RPMI + 10% FBS + 10ug/ml LPS and 25ng/ml of PMA or 1X RPMI + 10% FBS as a control. Cells were plated at a concentration of 10,000 cells/well in a volume of 200ml of media and cultured for 24 hours at 37°C with 5%CO2. After 24 hours cells were pelleted, and supernatant was removed and stored at -80°C for further analysis. IL-10 samples were diluted to a factor of 1:4 and ELISA assay was performed using manufacturer’s instructions (OptEIA™, BD Biosciences).

### Peritoneal cavity wash and Immunoglobulin quantitation with ELISA at postnatal day 14

Peritoneal washouts were performed by injecting 1mL of cold media into the PEC of P14 pups. Small piece of skin was removed on the lower left quadrant of the mouse PEC and needle was inserted through peritoneum. 1mL of media was injected and mice were shaken 10-15 times and media was pipetted out of the abdomen via a cut beneath the sternum. Contaminated samples either because of intestinal leakage and/or visible blood in the supernatant were discarded. Sample was filtered with 70uM filter and 500ul of sample was taken for cellularity counts via hemocytometer and supernatant analysis. Supernatant was stored at - 20°C until further analysis.

MaxiSorp plates were coated with 50ul Ig capture antibody (2ug/mL) or Phosphorylcholine-BSA (PC-BSA) (10ug/ml) for anti-PC IgM ELISAs and incubated for 2 hours at 37C. Plates were washed 3 times and blocked with 1X PBS + 0.5% Tween 20 + 0.5% BSA and incubated for 1 hour at RT. Plates were washed and samples and standards (2-fold serial dilution) were added and incubated for 2 hours at RT. Plates were washed and 50ul of appropriate detection (1:7000 dilution, or 1:2000 dilution for PC ELISA) was added to wells and incubated for 1 hour at RT. Plates were washed and 50ul TMB substrate was added to all wells and incubated for ∼15 mins at RT. 50ul of 1N sulfuric acid was added to stop the TMB reaction and plate was read at 450nm immediately following the addition of stop solution. PEC samples were normalized to cellularity (Total concentration/Total PEC cellularity).

### Quantification and Statistical Analysis

Prism v8.0 (GraphPad Software, San Diego, CA, USA) was used to perform statistical tests. Transplantation assays were randomized across age, sex, and cage for any given experiment. Investigator was blinded to treatment during preparation and analysis of samples. Statistically significant differences between groups with similar variance as determined by the SEM and SD were assessed by either un-paired two-tailed t-tests for parametric data, or Mann–Whitney tests for non-parametric data. 2-way ANOVA with Tukey’s multiple comparisons was used for comparison of GFP+ drHSC and Tom+ HSC under pIC or saline conditions. p values were calculated and are indicated as follows: *p ≤ 0.05; **p ≤ 0.01; ***p ≤ 0.001; ****p ≤ 0.0001. Sample sizes are indicated in each figure and unless otherwise noted, all experiments were performed in at least three independent replicates.

