## Supplemental Figures for "Prenatal inflammation perturbs fetal hematopoietic development and causes persistent changes to postnatal immunity"

**Supplementary Figure Legends**

**Supplementary Figure 1. The impact of prenatal inflammation induced by MIA on fetal and perinatal development and adult hematopoiesis**

**A)** Crown-rump length (CRL) of E15.5 fetuses following pIC or saline treatment. N=4-8 pups/litter representing at least 4 litters/condition.

**B)** Occurrence of gestation day at birth in pIC or saline treated litters. N=8-10 litters/condition.

**C-D)** Litter size **(C)** and weight **(D)** of mice shown in **B** on postnatal day (P)14.

**E-F)** Absolute cellularity of HSPCs **(E)** and CD45+ cells **(F)** at E15.5 in the fetal liver (FL) from pIC or saline treated mice shown in **Fig. 1C-D**.

**G)** Absolute cellularity of myeloid progenitors (GMPs, FcrIII/II+CD34+; CMPs, FcrIII/II, CD34+; MEPs, FcrIII/II-CD34-) in the FL at E15.5 in the same mice described in **Fig. 1C-D**.

**H)** Absolute cellularity of B-cells (CD45+CD19+B220+) in the FL at E15.5 in the same mice described in **Fig. 1C-D**.

**I)** Absolute cellularity of myeloid progenitors including MkP, GMP, pre-GM, PreMegE, and Pre-CFU-E, as defined by (Pronk et al., 2007) in the FL at E17.5 in pIC or saline treated litters. N= 6-7 representing at least two separate litters/experiments per condition.

**J)** Absolute cellularity of common lymphoid progenitors (CLPs; Lin-CD45+Ckit$^{lo}$Sca$^{lo}$IL7R+Flk2+) in the FL at E17.5 in pIC or saline treated litters. N= 13-24 representing at least three separate litters/experiments per condition.

**K)** Absolute cellularity of T-cells, B-cells, and granulocyte/macrophage (GM) cells in the FL at E17.5 in pIC or saline treated litters. N= 6-7 representing at least two separate litters/experiments per condition.

For all of the above experiments, data represents average $\pm$ SEM. ***p ≤ 0.001; Omission of statistical demarcation indicates differences are not statistically significant.

**Supplementary Figure 2. Impact of MIA on markers of inflammation in the fetal liver**

**A)** Representative FACS plots of Sca-1 expression in adult BM HSPCs one day after pIC or saline treatment.

**B)** Representative histogram of Sca-1 expression in adult LT-HSCs as shown in A.

**C)** Absolute quantification of FL Sca+1 cells in the same mice as described in **Fig. 1C-D**.

**D)** Representative FACS plots of Sca-1 expression in E15.5 FL CD45+ Lin- cells one day after pIC or saline treatment. Relative frequency of Sca-1+ cells is indicated.

**E)** Quantification of Sca1 MFI in FL LT-HSCs one day following treatment with pIC or saline.

**F)** Representative histograms of Sca-1 expression of LT-HSCs in FL one day following treatment with pIC or saline.

**G)** Heatmap displaying cytokines measured in maternal serum by LegendPlex multiplex assay at E15.5. N = 3 per condition.

**H)** Intracellular and extracellular TLR3 expression in fetal hematopoietic populations one hour following saline and pIC treatment. N = 8-14 representing 2-3 litters/condition. Data represent average $\pm$ SD, ****p ≤ 0.0001.

**I)** Single cell RNA-sequencing clusters scored against database of pathways (reactome.org) integral to interleukin signaling in the immune system. Scoring indicates enrichment assessed by the AddModuleScore function within individual clusters in each condition. Wilcoxon signed rank test. *p <0.05; ***p <0.001; ns, not significant.

**J, K)** Frequency of cells labeled by Tom+ fluorescence in myeloid progenitors (**J**) and B1-B cells (**K**) 1- or 24-hours post pIC or saline in Mx1-Cre pregnant mice. Labeling was determined by subtracting average labeling in saline treated mice from labeling in pIC-treated mice. N=6-11 representative of at least two independent experiments. Data represent average $\pm$ SD.

**L)** Representative FACS plots demonstrating gating strategy used to identify HSPC subsets derived from cultured GFP+ drHSCs or Tom+ HSCs 24 hours post-MIA with 20mg/kg pIC or saline.

**Supplementary Figure 3. Impact of MIA on fetal HSC function through transplantation**

**A)** Frequency of GM, B-cells, and T-cells within the reconstituted PB of recipients shown in **Fig. 5B-C.**

**B-E)** 18-week bone marrow (BM) chimerism in recipients of Tom+ HSCs or GFP+ drHSCs following saline or pIC as described in **Fig. 5B-C**. Chimerism in HSPCs, LT-HSCs, and ST-HSCs **(B)**; MPP2, MPP3, and MPP4 **(C);** granulocyte/macrophage progenitors (GMP), megakaryocyte progenitors (MkP), erythrocyte progenitors (EPs) and common lymphoid progenitors (CLP) **(D);** and granulocytes/macrophages (GM; CD11b+, Gr1+), B-cells (CD19+) and T-cells (CD3+) **(E)**. Data represent average $\pm$ SEM. $^*p \leq 0.05$; $^{**}p \leq 0.01$; $^{***}p \leq 0.001$; $^{****}p \leq 0.0001$.

**F)** Frequency of GM, B-cells, and T-cells within the reconstituted PB of secondary recipients shown in **Fig. 5D-E.**

**G-J)** 18-week bone marrow (BM) chimerism in secondary recipients of WBM from primary transplants of Tom+ HSCs or GFP+ drHSCs following saline or pIC described in **Fig. 5D-E.** Chimerism in HSPCs, LT-HSCs, and ST-HSCs **(G)**; MPP2, MPP3, and MPP4 **(H)** granulocyte/macrophage progenitors (GMP), megakaryocyte progenitors (MkP), erythrocyte progenitors (EPs) and common lymphoid progenitors (CLP) **(I);** and granulocytes/macrophages (GM; CD11b+, Gr1+), B-cells (CD19+) and T-cells (CD3+) **(J)**. Data represent average $\pm$ SEM. $^*p \leq 0.05$; $^{**}p \leq 0.01$; $^{***}p \leq 0.001$; $^{****}p \leq 0.0001$.

**Supplementary Figure 4. Molecular impact of MIA on fetal HSPCs**

**A)** Heat map of top seven most differentially expressed genes per cluster following pIC or saline treatment. Legend represents normalized gene expression. Plot generated using DoHeatmap.

**B)** UMAP plot depicting cell-cycle states (G0/G1, S, G2/M) of every cell across all 14 clusters based on Cyclone scoring.

**C)** UMAP plots depicting enrichment of eGFP or tdTomato transcripts across all 14 clusters in response to pIC or saline. Color bar indicates the enrichment scoring.

**D-E)** Violin plot showing relative expression of marker genes associated with C2 across all clusters, including *Mllt3* **(D)** and *Meis1* **(E).** Wilcoxon signed rank test with respect to background. $^{***}p \leq 0.001$.

**F-G)** Violin plots showing scoring of megakaryocyte **(F)** and myeloid **(G)** identity across all clusters (Cabezas-Wallscheid et al., 2014). Wilcoxon signed rank test with respect to background. $^*p \leq 0.05$; $^{***}p \leq 0.001$.

**H-K)** Violin plot showing relative expression of marker genes associated with C9 across all clusters including *Spi1* **(H)**, *Pbx1* **(I)**, *Cd74* **(J)** and Tet2 **(K)**. Wilcoxon signed rank test with respect to background. *$p \leq 0.05$; **$p \leq 0.01$; ***$p \leq 0.001$.

**Supplementary Figure 5. Prenatal inflammation influences postnatal hematopoiesis**

**A-E)** Absolute quantification of adult HSPCs for mice shown in **Fig. 6K-M, O**. N = 6 representing at least three separate litters/experiments per condition. Data represent average ± SEM.

**F)** Frequency of adult BM HSPCs for mice shown in **SFig. 6E.**

**SFigure 1. The impact of prenatal inflammation induced by MIA on fetal and perinatal development.**

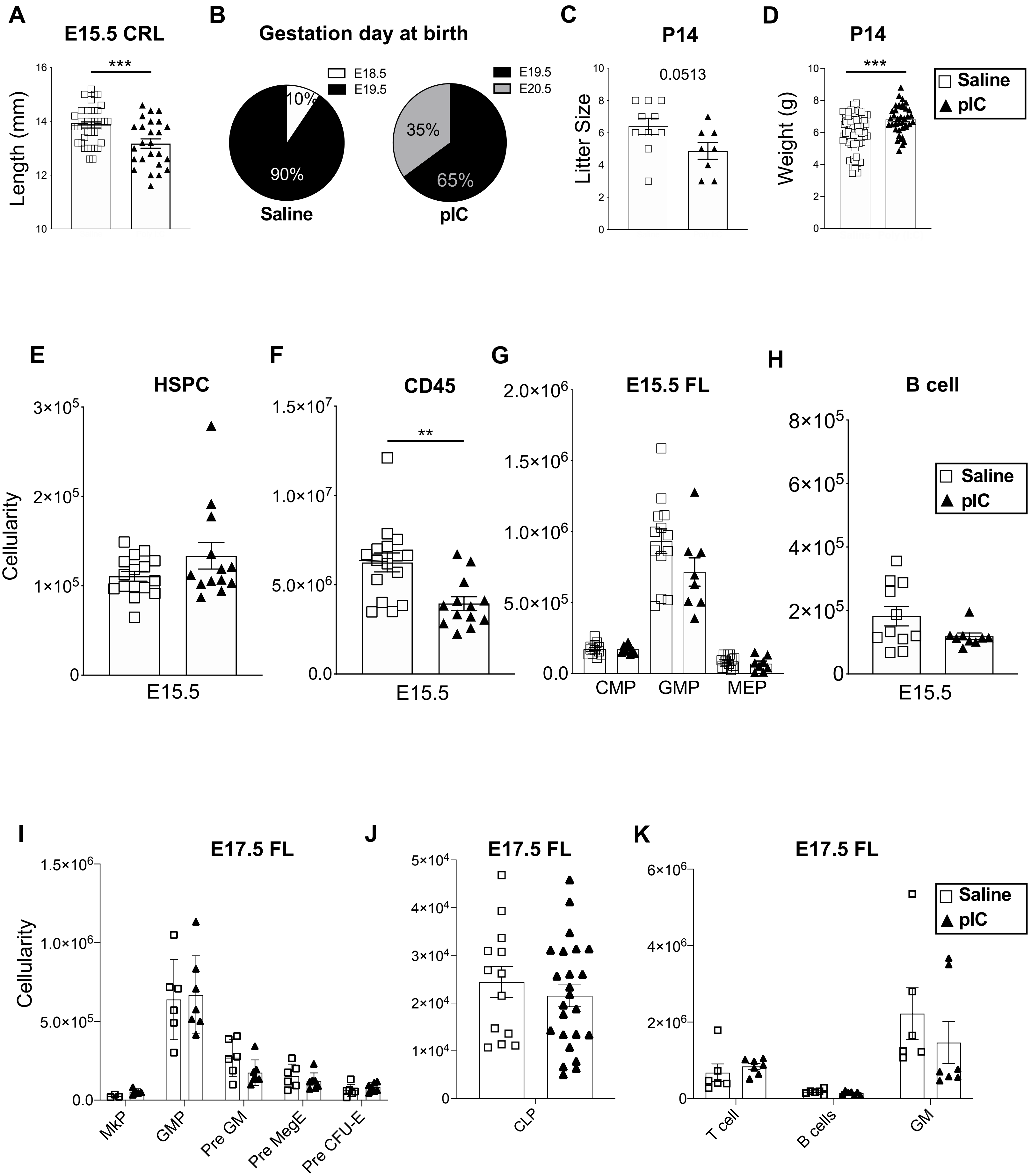

### Sfigure 2. Impact of MIA on markers of inflammation in the fetal liver

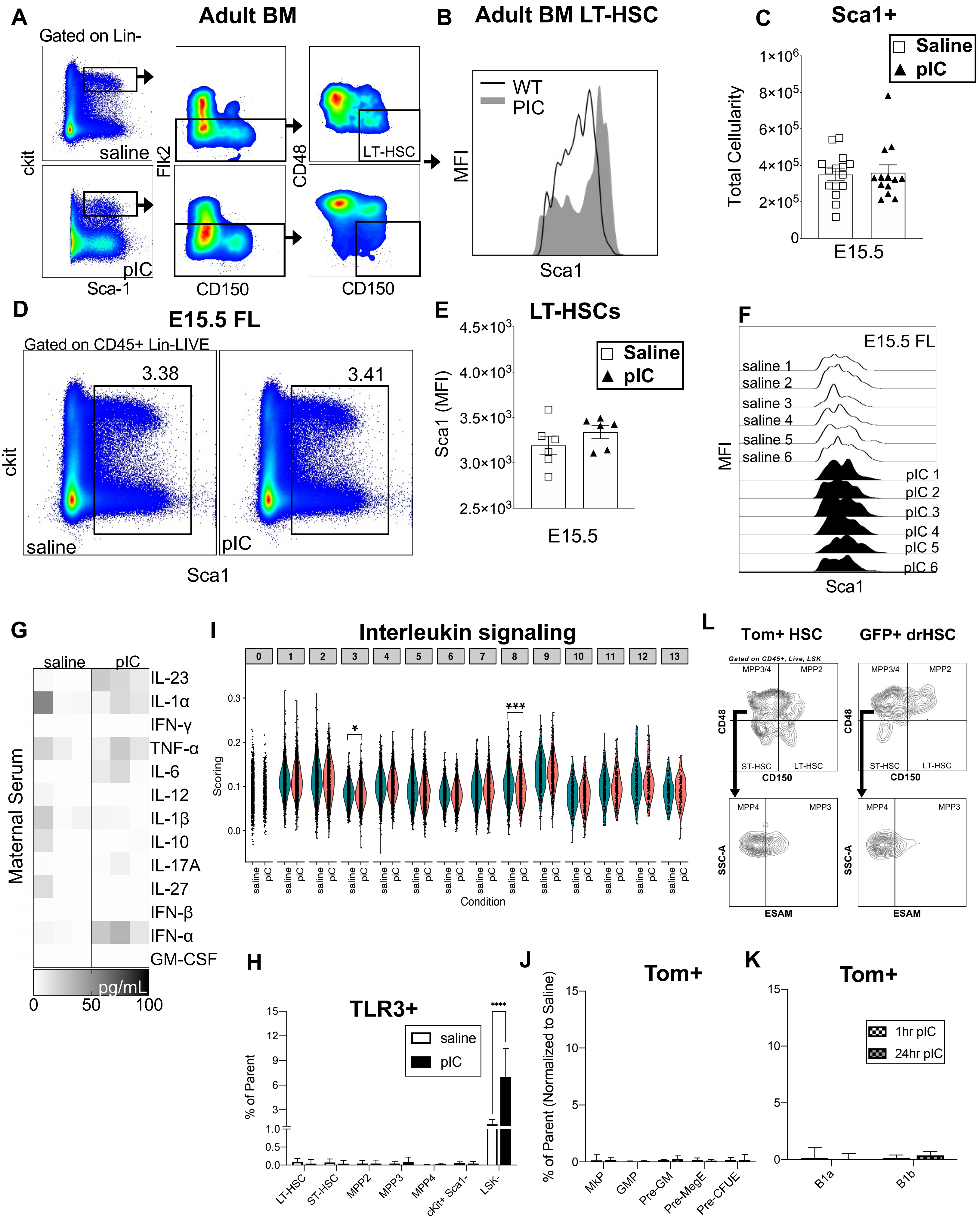

### SFigure 3. Impact of MIA on fetal HSC function through transplantation

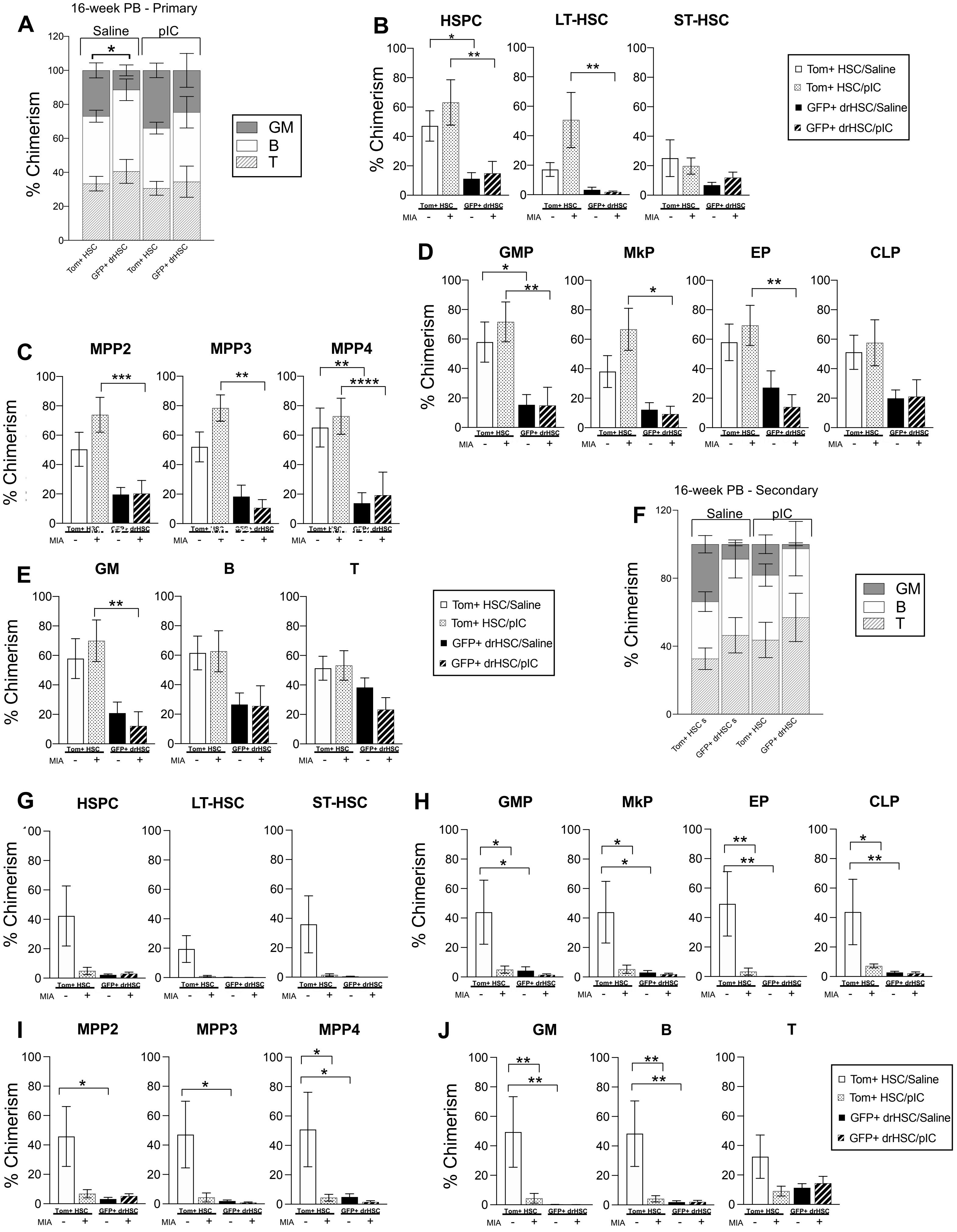

### Sfigure 4. Molecular impact of MIA on fetal HSPCs

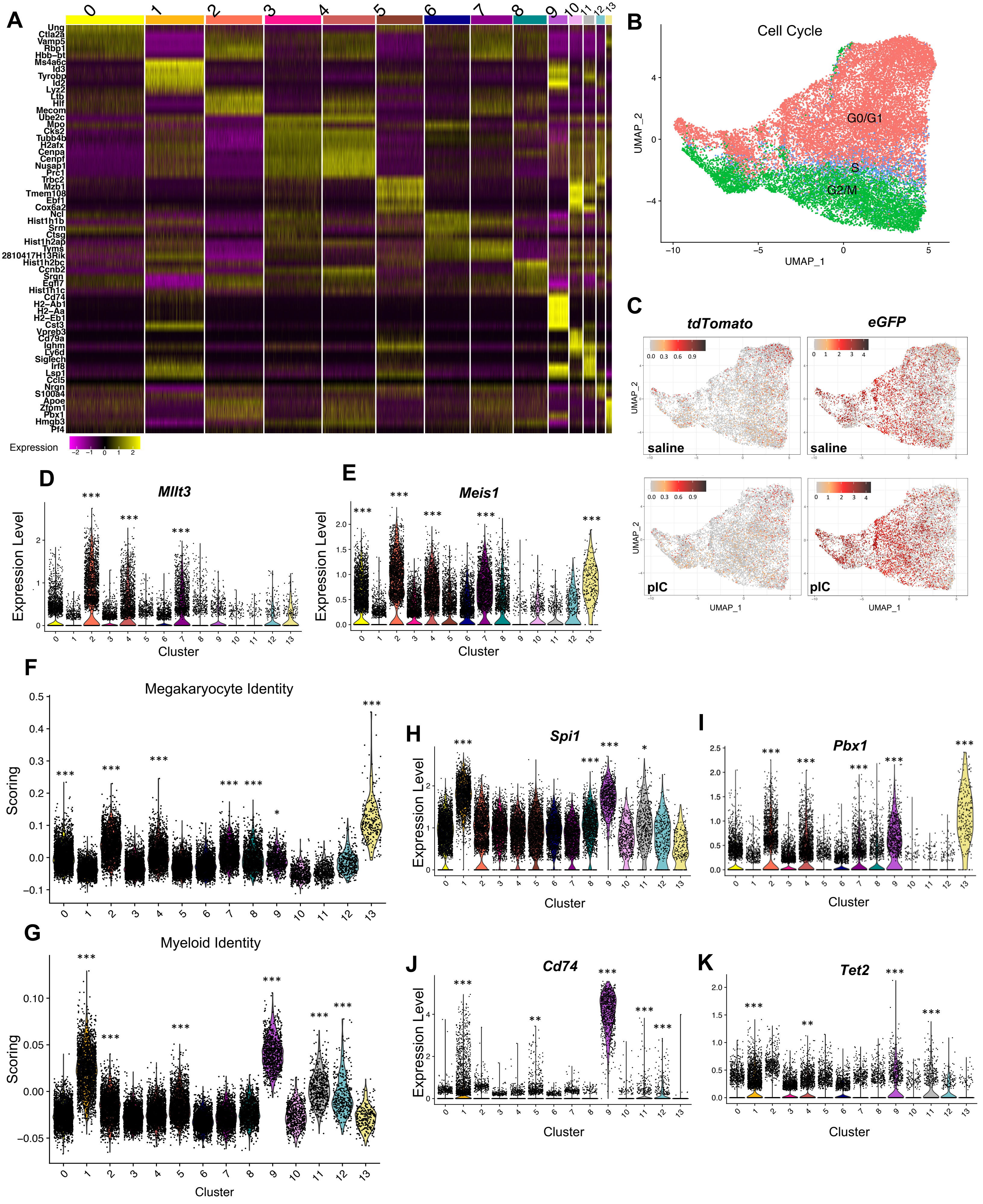

**Sfigure 5. Prenatal but not adult inflammation influences postnatal hematopoiesis**

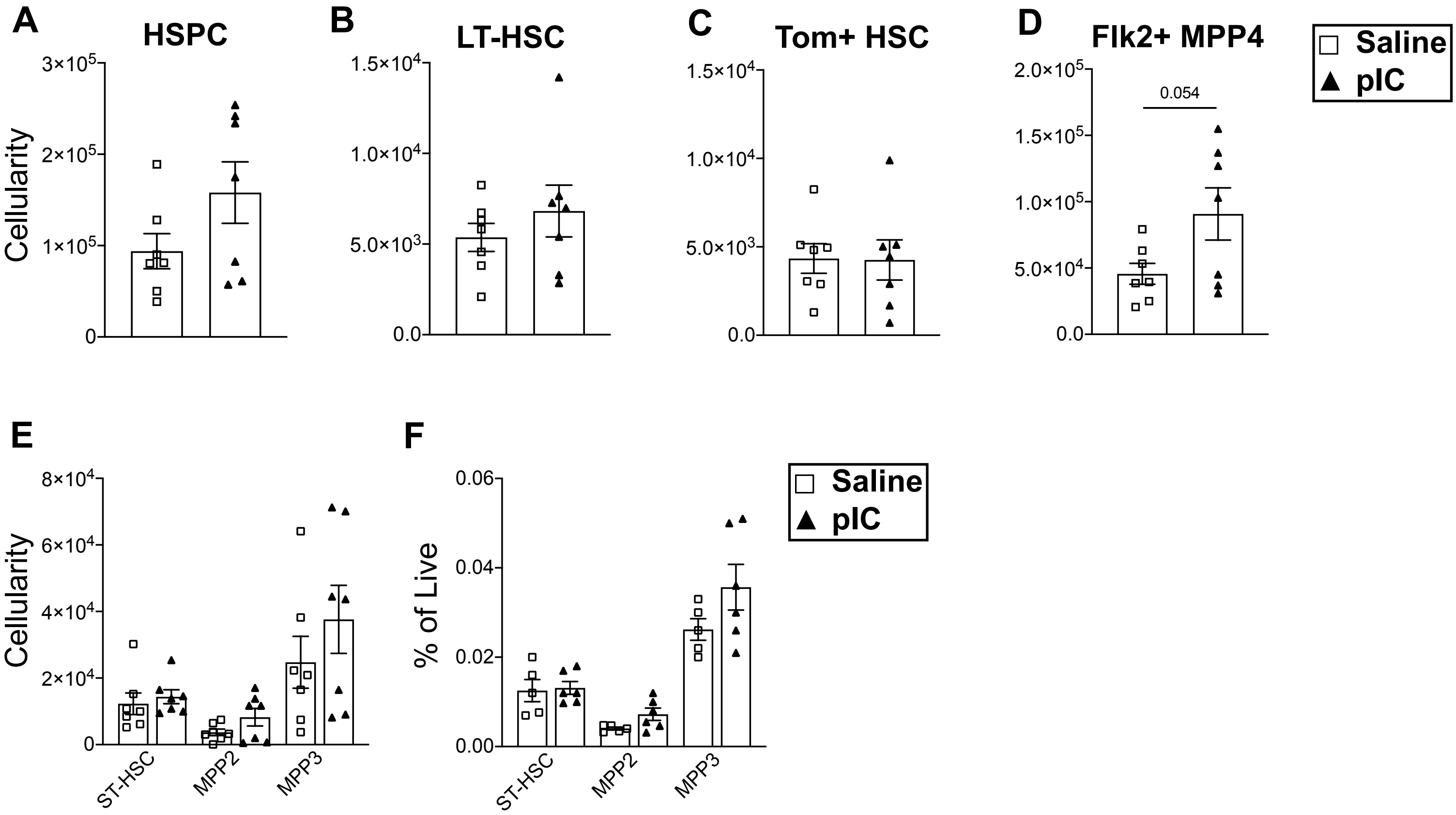
